# Nanocoding: Lipid Nanoparticle Barcoding for Multiplexed Single-Cell RNA Sequencing

**DOI:** 10.1101/2024.12.16.628827

**Authors:** Yujun Feng, Donglai Chen, Catherine C. Applegate, Natalia Y. Gonzalez Medina, Chia-wei Kuo, Opeyemi H. Arogundade, Chris L. Wright, Fangxiu Xu, Jenny Drnevich, Andrew M. Smith

## Abstract

Sample multiplexing is an emerging method in single-cell RNA sequencing (scRNA-seq) that addresses high costs and batch effects. Current multiplexing schemes use DNA labels to barcode cell samples but are limited in their stability and extent of labeling across heterogeneous cell populations. Here, we introduce Nanocoding using lipid nanoparticles (LNPs) for high barcode labeling density in multiplexed scRNA-seq. LNPs reduce dependencies on cell surface labeling mechanisms due to multiple controllable means of cell uptake, amplifying barcode loading 10-100-fold and allowing both protection and efficient release by dissolution. In cultured cell lines and heterogeneous cells from tissue digests, Nanocoding occurs in 40 minutes with stability after sample mixing and requires only commercially available reagents without novel chemical modifications. In spleen digests, 6-plex barcoded samples show minimal unlabeled cells, with all barcodes giving bimodal count distributions. Challenging samples containing lipid-rich debris and heterogeneous cells from adipose tissue of obese rodents show more than 95% labeling with all known subtypes identified. Using Nanocoding, we investigate gene expression changes related to aging in adipose tissue, profiling cells that could not be readily identified with current direct conjugate methods using lipid or antibody conjugates. This ease of generating and tuning these constructs may afford efficient and robust whole-sample multiplexing with minimal sample crosstalk.

---

Single-cell RNA sequencing (scRNA-seq) has recently become a widely used method for cell transcriptomic profiling, divulging extensive insights into cell heterogeneity,^1^ with adoption across communities in the life and medical sciences.^2^ The past decade has seen a surge in the volume of information obtainable from single cells using scRNA-seq driven by advances in microfluidic and microwell devices as well as bioinformatics tools, now achieving ∼10,000 cell reads as ∼3 terabyte data sets.^3,4^ A current technological bottleneck is that scRNA-seq methods apply single-use, single-sample devices, setting cost barriers and leading to batch effects when comparing across samples.^5^ While next-generation devices will increase both sample number and cell reads, measuring multiple samples on single devices is needed to address these limitations.^6–9^

Incorporating multiple samples into a single library preparation (sample multiplexing) is becoming practical for single cells in a similar way as for multiplexed bulk DNA sequencing.^8–11^ The strategy applies exogeneous single-stranded DNAs (ssDNAs) as sample-specific identifiers (barcodes) before mixing samples so that the sample origin can be read by sequencing within a single library preparation.^11^ The most common methods apply ssDNA conjugates of either antibodies that target cell surface antigens or lipids that associate with the plasma membrane, both of which efficiently label certain samples like peripheral blood mononuclear cells (PBMCs).^11–13^ Accurate identification of the sample origin of each cell from bioinformatic post-processing requires a distinct bimodal distribution of barcode counts across the cells in the mixed sample so that thresholds distinguishing labeled cells are unambiguous.^11^ As such, barcoding requires high-density labeling, stability of the barcode, lack of transfer between cells in the mixed sample, and efficient release and incorporation into sequencing machinery to minimize unidentifiable cells and incorrect sample assignments.^10,14^

Current barcoding methods with antibodies and lipids are problematic for samples with heterogeneous cell types that are variable in antigen expression and their interactions with membrane-adsorbing labels.^14^ Both label types function by reversible interactions that depend on environmental conditions, washing steps, and dilution factors, which pose challenges in the workup and numerous processing steps between labeling and sequencing.^15,16^ Further, in both droplet and microwell-based scRNA-seq devices, each cell capture compartment is larger than the cell, which can result in background deriving from free RNA or dissociated barcodes in the ambient solution (**Fig. 1a**).^17,18^ Antibody-based labeling is further limited by the lack of a single antigen expressed at high levels across all cells in a population. Recent comparisons of lipid-ssDNA barcodes across samples and lipid types showed variable performance without a clear molecular explanation or generalizable protocol,^10,14^ and label ‘crosstalk’ (from 2% to 15%) has been reported after sample mixing for 20 minutes on ice.^14^ Recommended approaches, therefore, include pilot tests to screen for optimal label types or bioinformatic processing by presuming barcode assignment through statistical hypotheses built upon sample-variant labeling,^19^ but inconsistent results may still be expected for non-specialists. Novel labeling approaches include intracellular delivery by transient transfection and DNA ligation in permeabilized cells,^20–22^ methods that could alter RNA levels, lead to variable barcoding efficiency, or cause toxicity. A more universal plasma membrane labeling strategy uses active esters such as *N*-hydroxysuccinimide (NHS) to ligate ssDNA barcodes to ε-amines of lysine residues or N-terminal amines of membrane proteins.^23,24^ However the amine targets are also present on soluble proteins and tissue debris, and active esters can have variable reactivity, leading to low barcode counts or non-bimodal barcode distributions compared with antibody-based conjugates in PBMCs.^23,25^ At present, a stable and sample-type independent strategy is still not available for multiplexed scRNA-seq.^10^

**Fig. 1.**
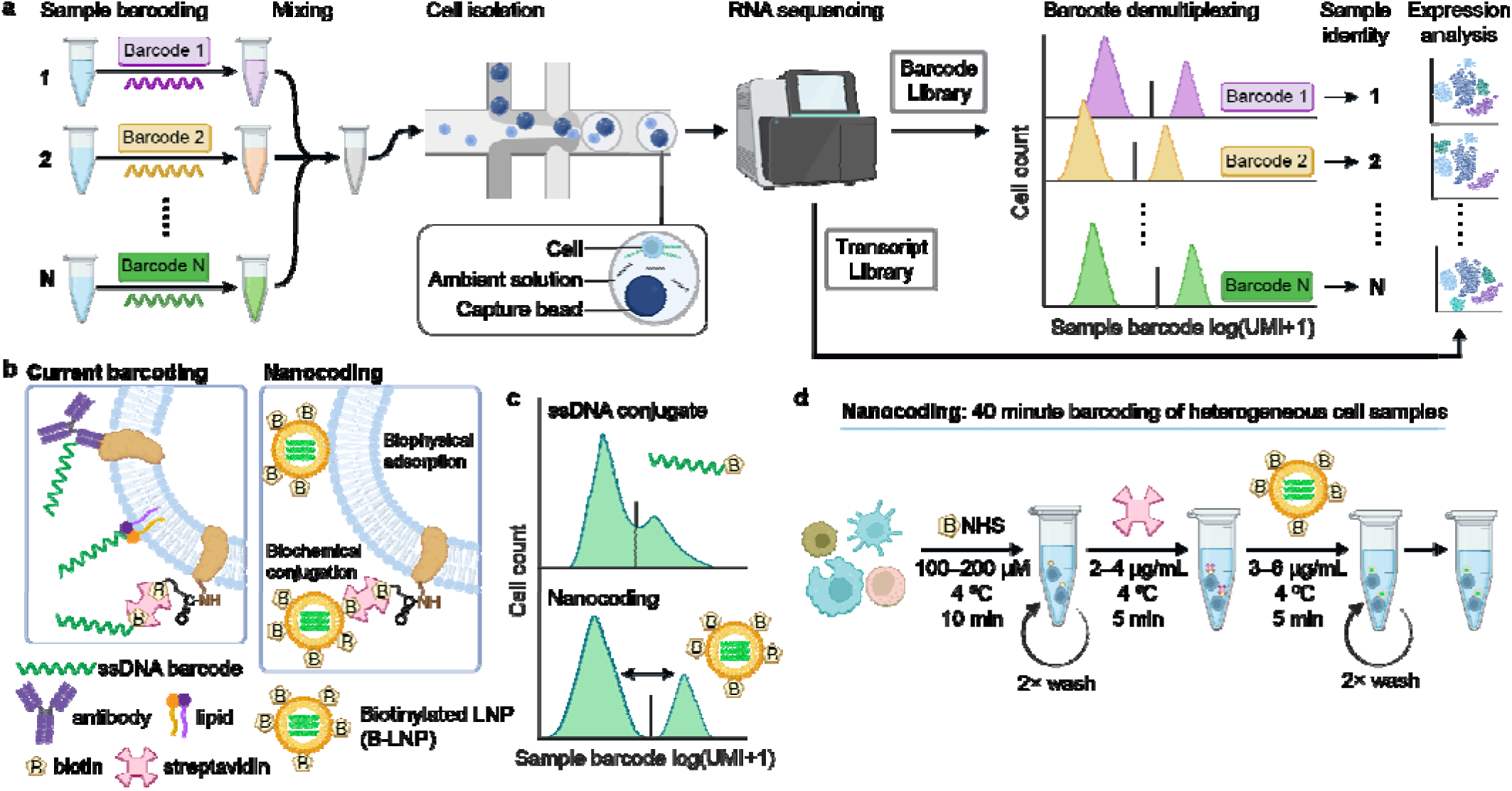
Nanocoding design and workflow for sample multiplexing in single-cell RNA sequencing (scRNA-seq). **(a)** Workflow shows barcoding of N cell samples, sample mixing, microfluidic single-cell isolation, and RNA sequencing to generate barcode and RNA transcript libraries. The barcodes are detected in bimodal distributions across the cells to allow demultiplexing of each cell’s sample, yielding sample-specific cell expression profiles from the transcript library. **(b)** Comparison between Nanocoding and current methods to barcode cells, including covalent ssDNA barcode conjugates of antibodies targeting cell surface antigens and lipids that adsorb to plasma membranes. Cell surface proteins can also be biotinylated using biotin-NHS reagents that bind to tetravalent streptavidin (SAv) which binds to biotinylated ssDNA. Nanocoding functions through a combination of biophysical adsorption and biochemical conjugation, using biotin-functionalized lipid nanoparticles (B-LNP) encapsulating ssDNA barcodes. **(c)** Example of improvement in barcode distribution for multiplexed scRNA-seq using Nanocoding (bottom) relative to biotinylated ssDNA (top). **(d)** Workflow for Nanocoding labeling of heterogeneous cells showing cell surface biotinylation, SAv binding, and B-LNP binding.

In this work, we develop “Nanocoding” with lipid nanoparticles (LNPs) to deliver a large number of oligonucleotides to diverse cells for multiplexed scRNA-seq (**Fig. 1a**). Unlike current labels deriving from chemical biology and molecular biology tools with 1:1 stoichiometries between anchoring moieties and barcodes, LNPs are a class of nanomaterials allowing high-density encapsulation of hundreds of ssDNA strands into single colloidal entities through self-assembly (**Fig. 1b**).^26,27^ LNPs are broadly applied in healthcare as a nucleic acid delivery vehicle with minimal cytotoxicity due to extensively developed lipid chemistries. Materials and formulation methods are well developed and require only common lab instruments, and do not require custom synthesis or expensive chemical modifications for nucleic acids.^28,29^ ^30^ Importantly, diverse routes of rapid, irreversible cell association are enabled by selecting from diverse lipid libraries, including surface tethering of biochemical ligating groups (*e.g.*, biotin) and tunable cell endocytotic processes driven by biophysical adsorption.^31^ As we show here, a mixed cell population can be comprehensively labeled through a combination of these modalities, while barcodes remain protected within LNPs with efficient release upon cell lysis. We optimize methods for concurrent labeling by biophysical adsorption and irreversible labeling through surface amine-targeted NHS linkers followed by streptavidin and biotin-functionalized barcode-encapsulated LNPs. We evaluate Nanocoding in primary and cultured cells as well as stromal vascular fraction cells from digests of mouse adipose tissue, a lipid-laden, heterogeneous sample with a high fraction of CD45^−^ and CD45^lo^ populations, which can be challenging for existing methods. We apply Nanocoding to compare features from adipose tissue cells from aged and young mice in the state of obesity, profiling age-related inflammatory features and lipid-transfer activity in macrophages, T cells, and non-immune cells.

## RESULTS AND DISCUSSION

### Nanocoding LNP Design and Evaluation

Oligonucleotides used for scRNA-seq barcoding are 90-mer ssDNA sequences designed for 10X Genomics platforms (**Table S1**). We evaluated a breadth of LNP formulations to determine capacities for biophysical and chemical labeling of live cells (**Extended Fig. 1**, **Table S2**).^32^ LNPs were designed using a fixed base formulation (**Table S3**) including a methoxy-terminated polyethylene glycol (PEG; 2,000 Da) functional lipid (PEG-DMG).^28^ PEG-DMG could be substituted with a similar lipid functionalized with biotin (B-PEG-DSPE) to generate a biotinylated LNP (B-LNP) that irreversible chemically ligates to cell surfaces when applied in combination with NHS-functionalized biotin linkers (B-NHS) and streptavidin (SAv) (**Fig. 1b**), reagents broadly applied in molecular biology and targeted delivery.^33^ The base B-LNP formulation also included the ionizable lipid DLin-MC3-DMA (MC3), cholesterol, and DSPC in ratios optimized for stability, dense encapsulation of nucleic acids, and efficient gene delivery to cells.^27,28^

By tuning the ionizable or non-ionizable cationic lipid responsible for condensing ssDNA in the LNP core, the zeta potential could be controlled to be moderately cationic or anionic (–4 to +3 mV) in neutral solution while the sizes ranged between 50–120 nm (**Extended Fig. 1a-d**). The MC3 lipid allowed the highest encapsulation of ssDNA barcodes per LNP compared with other ionizable lipids as well as the highest capacity for labeling by both biophysical adsorption and B-NHS-specific association by flow cytometry (**Extended Fig. 1e-f**). Notably, the lipid ALC-0315 nearly eliminated biophysical adsorption and almost exclusively facilitated chemically specific labeling. In contrast, the cationic lipid resulted in exclusively biophysical labeling (**Extended Fig. 1e**) likely due to a slightly cationic surface charge (+3 mV) which can lead to cell toxicity.^34,35^

The lead candidate B-LNPs based on MC3 had a diameter of 90 ± 3.2 nm (mean ± s.e.m.) by cryo-electron microscopy (**Fig. 2a**) and 100 nm by dynamic light scattering (PDI 0.11, **Fig. 2b**), with zeta potential of –2.2 mV. These parameters, as well as ssDNA encapsulation efficiency (**Table S4**), are in the range of values previously described for similar formulations.^36^ The B-LNP hydrodynamic diameter was dependent on the lipid nitrogen-to-DNA phosphate (N:P) ratio, with size slightly larger at N:P=3:1 (**Fig. 2c**) which also led to the highest barcode encapsulation number (85) by fluorescence correlation spectroscopy (FCS), both per particle (**Fig. 2d**) and by particle volume (**Fig. S1**). This encapsulation efficiency is at the high end of expectations (10–100) from estimates of 2–5 copies of 996-mer mRNA.^37^ For this LNP, SAv binding per LNP could be tuned between 2–40 by adjusting the amount of B-PEG-DSPE between 0–3% (**Fig. 2e**), close to estimates of 80 per particle for 50 nm unilamellar liposomes.^38^ In all following experiments, we used the formulation of 3:1 for N:P with 1% B-PEG-DSPE to enhance labeling through multivalency.

**Fig. 2.**
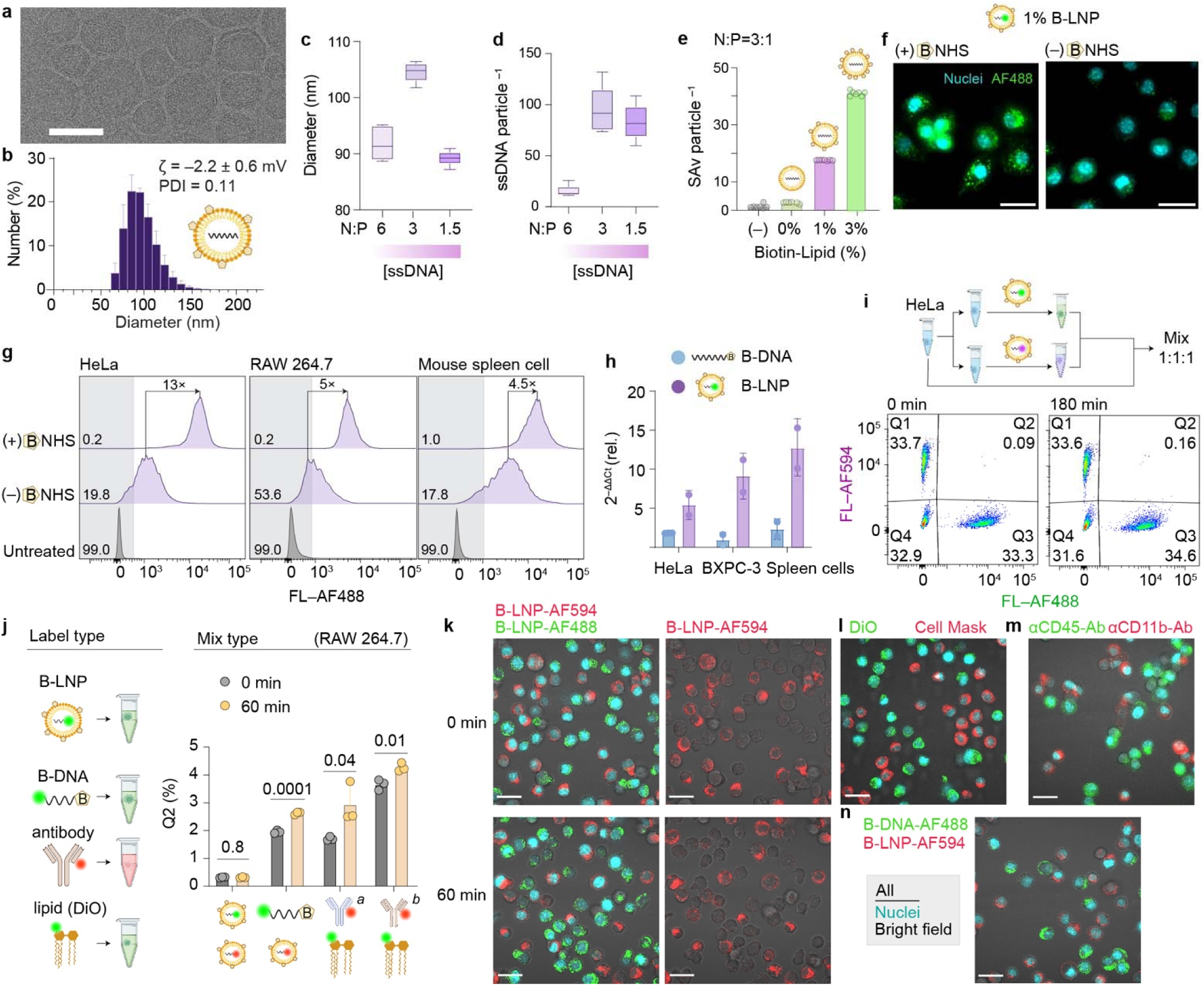
Characterization of biotinylated LNPs (B-LNP) and Nanocoding workflow in live cells. **(a)** Cryo-electron micrograph of B-LNP with 1% Biotin-PEG-DSPE (Biotin-Lipid) encapsulating 90-mer ssDNA barcode (**Table S1**) at 3:1 nitrogen-to-phosphate (N:P). Scale bar, 100 nm. **(b)** B-LNP hydrodynamic diameter by dynamic light scattering (DLS) including polydispersity index (PDI) and zeta potential (ζ). **(c)** DLS-derived Z-average mean of B-LNP with indicated N:P values. **(d)** Mean number of ssDNA per B-LNP particle based on ssDNA encapsulation assay and LNP concentration from fluorescence correlation spectroscopy. **(e)** Mean number of binding streptavidin (SAv) per LNP particle at indicated Biotin-Lipid composition. **(f)** Fluorescence micrographs of RAW 264.7 cells treated with B-NHS, SAv, and B-LNP containing AF488-DNA (**Table S1**). On the right, the treatment was the same except B-NHS was excluded. Scale bar: 20 μm. **(g)** B-LNP labeling of cultured and primary mouse spleen cells treated with or without B-NHS. Differences in median fluorescence are shown together with the percentage of unlabeled cells from flow cytometry. **(h)** Cell labeling by PCR of DNA extracts from cell lysate, which includes barcodes from both cell-labeled and the ambient solution, to compare biotin-ssDNA conjugates (B-DNA) and B-LNP. For each type of label, the 2^−ΔΔCt^ value reflects the PCR cycle number for B-NHS treated cells relative to cells not treated with B-NHS. *N*=2 independent biological replicates. Details are in **Methods**. **(i)** B-LNP label stability in mixed samples of HeLa cell. Gates Q1–4 were set using cell samples labeled with only one B-LNP color. The percentage of cells in each gate is shown. **(j)** Label stability in mixed samples of RAW 264.7 cells using combinations of B-LNP, B-DNA, antibody, and lipid (DiO) labels. The Q2 percentage of the population is shown immediately after mixing and 60 minutes later. *N*=3 independent biological replicates. *a,* αCD45 antibody, *b*, αCD11b antibody. Data represent mean ± s.d. *p*-values are determined by two-tailed unpaired Student’s *t*-test. Raw data are shown in **Figures S6**, **S7**, and **S8. (k)** Fluorescence micrographs of RAW 264.7 cells that labeled with either B-LNP-AF488 (green) or B-LNP-AF594 (red) and then mixed. Nuclear stain (blue) and bright field (grayscale) are shown as an overlayed. Scale bar: 20 μm. **(l)** Same as k except cells that were labeled with either DiO (green) or CellMask Orange (red). **(m)** Same as k except cells that were labeled with either αCD45 antibody (green) or αCD11b antibody (red). **(n)** Same as k except cells that were labeled with either B-DNA (green) or B-LNP (red).

Based on fluorescence from dye-labeled oligonucleotides (3–6 μg/mL), B-LNP densely and comprehensively labeled a variety of cells treated with B-NHS and SAv at doses used in drug delivery to minimize toxicity and immunogenicity.^28^ This included cultured cell lines with large sizes (HeLa) and small sizes (RAW 264.7) and cells from mouse spleen digests (**Fig. 2f, 2g**). While each cell type was labeled by a combination of biophysical adsorption and chemically specific binding, labeling density was dominated by the latter, yielding a 4.5–13-fold median signal enhancement. Setting a specificity threshold of 99% by fluorescence intensity of unlabeled samples, >99% of the cells were distinguished as labeled in each cell type (**Fig. 2g**). Labeling density was enhanced when using B-NHS with a more flexible linker (**Fig. S2a**) indicating that steric hindrance may limit labeling.

### Comparison with Chemical and Biomolecular Labels

Labeling density and barcode specificity of B-LNP were compared with other cell surface labeling methods. A dye-labeled ssDNA barcode with terminal biotin (B-DNA, **Table S1**) could similarly label cells as with B-LNP after irreversible B-NHS and SAv attachment and could likewise efficiently label some cell types (**Fig. S3** and **S4**). However, labeling with B-DNA was not comprehensive for some cell types and was lower than for B-LNP on average based on flow cytometry (**Fig. S3b** and **Fig. S5**). Moreover, by qPCR of cell lysate, B-LNP yielded 5–10-fold higher 2^−ΔΔCt^ value of barcode DNA compared with B-DNA, suggesting an enhanced capacity of B-LNP to protect and release ssDNA barcodes for ensuing enzymatic processes (**Fig. 2h**).

To compare label stability with reversibly bound barcode labels, we used fluorescently labeled antibodies against surface antigens (CD45 or CD11b) and membrane dyes (DiO and CellMask Orange) and mixed samples to evaluate crosstalk by transfer of fluorescent signals. B-LNP showed negligible cross-labeled cells 1–3 hours after sample mixing (**Fig. 2i**) in both HeLa (0.16%) and RAW 264.7 (0.3%) cell lines (**Fig. 2j**). In contrast, DiO and antibodies showed higher cross-labeled populations (4–6%) immediately upon mixing with a small but statistically significant increase (0.5–1%) in cross-labeled cells after one hour which did not occur for B-LNP (**Fig. 2j**, **2k, Fig. S6, S7**). Surprisingly, cross-labeling was also observed for B-DNA barcodes bound through irreversible SAv interactions (**Fig. 2l**, **Fig. S8**). The difference in crosstalk between B-LNP and B-DNA may be due to the additional protection of ssDNA tags within LNPs compared with those that are attached “bare” to the cell surface.^31,39^

To understand the crosstalk resistance afforded by B-LNP labels, cells were labeled with B-LNPs containing ssDNA conjugates of dyes with distinct colors, then mixed and imaged live, maintaining the cells at 4 °C until imaging. Most LNPs were intracellular less than 10 min after sample mixing (**Fig. 2k**). This contrasts with B-DNA labels for which much remained located on the cell membrane with some cells appearing to be completely unlabeled (**Fig. 2l**, **Fig. S4**). Significant internalization was also evident for antibody and lipid-based stains (**Fig. 2m**, **2n**), however, there was still significant cell transfer over time of αCD45 antibodies (**Fig. S9**) and one membrane label (Cell Mask), which labels cell surfaces with slower internalization compared with DiO (**Fig. S10**)^40,41^ Similar trends in antibody label transfer were observed by flow cytometry (**Fig. S6** and **S7**). It is further important to emphasize that fluorescent analogs are expected to underestimate the extent of crosstalk relative to oligonucleotide labels, as the labeling affinity and stability are expected to be compromised by nucleic acid modifications.^30,42^

### Six-Sample scRNA-seq Multiplexing with Spleen Tissue Digests

Nanocoding was applied for scRNA-seq using a set of 6 barcodes (**Table S1**), each separately packaged into B-LNPs with 3 μg/mL ssDNA. These B-LNPs were used to barcode heterogeneous cell populations from fresh digests of spleens from C57BL/6J mice fed a low-fat diet (“lean,” *N*=3) or high-fat diet (“obese,” *N*=3) (**Fig. 3a**). All six samples showed high viability (>90% cells) after Nanocoding without live cell enrichment. The six samples were mixed in equal cell numbers and a total of 9000 cells were sequenced, targeting 1500 cells for each sample. Measured barcode counts by scRNA-seq exhibited bimodal cell count distributions across the cell mixture (**Fig. 3b**) and barcode counts were high using HTOdemux with cell numbers consistent with their input ratios (**Fig. 3b**).^11^ The number of multiplets (8%) was close to the expected value using this cell number and microfluidic chip (∼6.4% at 8,000 cells recovered for 10K library preparation kit). There were minimal identified cells that were unlabeled (<0.5%) with no apparent dependence on cell type (**Fig. S11**). Two commonly used methods to demultiplex applying different fitting or statistical methods (HTOdemux and deMULTIplex) showed minimal discordance in single cell, unlabeled cells, or multiplet assignments based on CellhashR (**Fig. 3c, Fig. S12**).^19^ The 6 cell samples separated well in *t*SNE plots by barcode number (**Fig. 3d).**

**Fig. 3.**
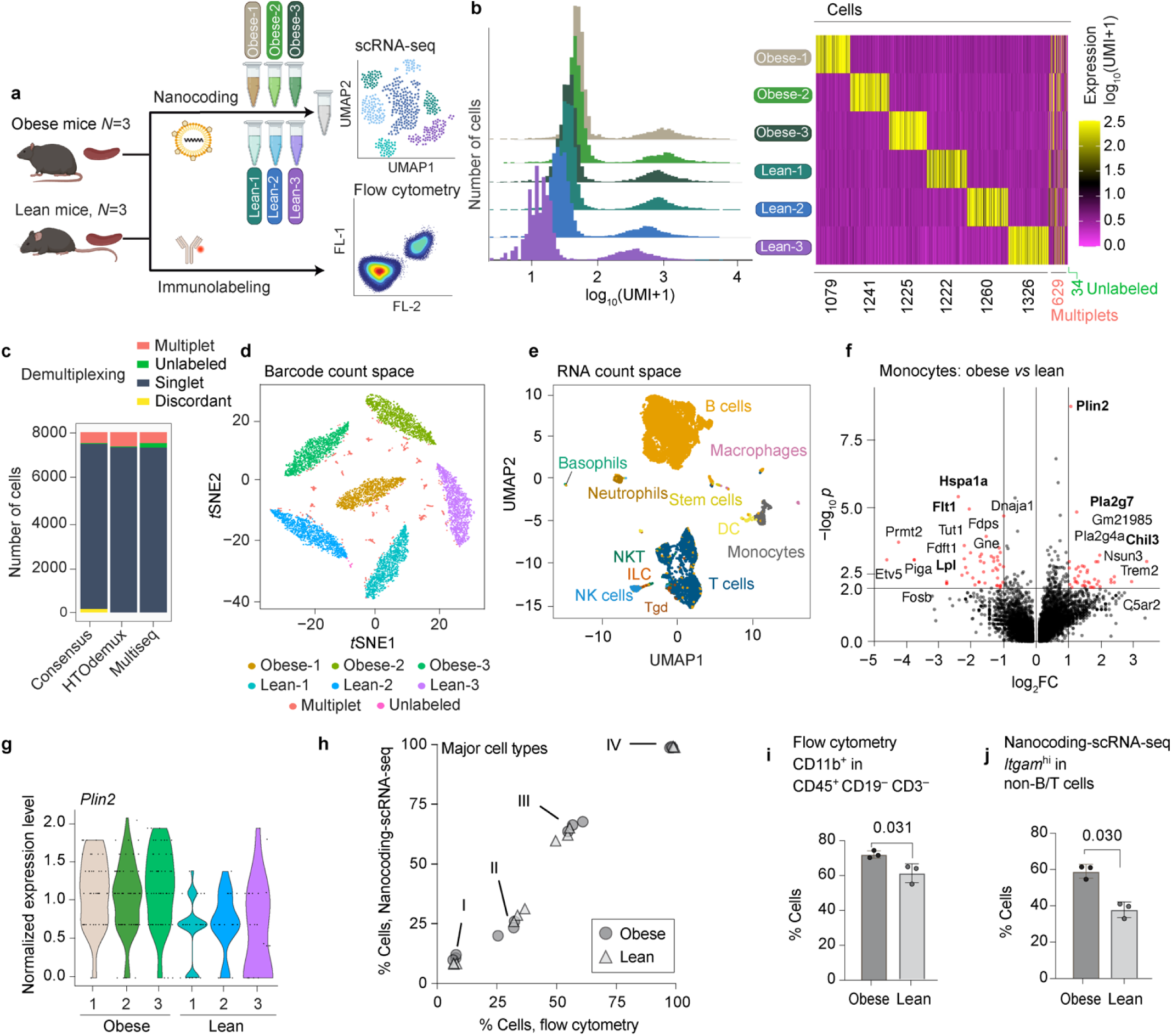
Multiplexed barcoding of mouse spleen digests by Nanocoding. **(a)** Experimental design of a six-sample spleen study using Nanocoding-based scRNA-seq. Spleens were from lean or obese mice, with cell populations comparisons by flow cytometry of the same samples. No live-cell enrichment was included. **(b)** Heatmap of log_10_(UMI+1) reads of 6 barcode sequences (6 rows) in 8019 cells (thin columns). Cells were sorted based on barcode identification using HTOdemux. **(c)** Comparison of cell identification results between HTOdemux and deMULTIplex based on barcode matrix only (CellhashR package). **(d)** Cell clustering in barcode count space as a *t*SNE plot, color-coded based on identified sample origin (CellhashR package). **(e)** Cell clustering in RNA expression space. Cell types are annotated automatically using the SingleR package with reference to the ImmGen database. **(f)** Differential gene expression of monocytes comparing obese *versus* lean mice, aggregating all samples within each biological condition. Selected top variated genes are labeled. Red dots have more than 2-fold change (FC) and *p* < 0.01 by single-cell analysis. Bold font indicates FC > 2 and *p* < 0.001 by pseudobulk analysis. **(g)** *Plin2* expression in monocytes in each sample with gene counts log-transformed and normalized (SCTransform function in Seurat package). **(h)** Comparison of major cell populations identified by multiplexed Nanocoding scRNA-seq or flow cytometry (FC). Cells are indicated as: I, CD45^+^ CD3^−^ CD19^−^ (FC) or myeloid lineage (scRNA-seq); II, CD45^+^ CD3^+^ (FC) or T cells (scRNA-seq); III, CD45^+^ CD19^+^ (FC) or B cell (scRNA-seq); IV, CD45^+^ (FC) or immune cells (scRNA-seq). (**i**) Percentage of CD11b^+^ cells in CD45^+^ CD3^−^ CD19^−^ cells (FC). (**j**) Percentage of *Itgam*^hi^ cells in non-B/T/NK cells (scRNA-seq). In **i** and **j**, data show mean ± s.d. with statistical significance determined by two-tailed unpaired *t*-test. Sequencing quality reports are summarized in **Table S6.** Bioinformatic analysis is included in **Methods** and **Code.** Six-plex data are in the **Data** section.

To verify cell numbers and sample barcode assignments, we compared the ratio of major immune cells between scRNA-seq and flow cytometry data sets from the same tissues tested on the same day. Immune cell types from scRNA-seq were annotated using SingleR with reference to the ImmGen database,^43,44^ with multiplets and unlabeled cells filtered, resulting in the primary immune cells expected for mouse spleen (**Fig. 3e**). The results were consistent in that the ratio of major immune cell populations was similar for multiplexed scRNA-seq and flow-cytometry tests (**Fig. 3h**). The results were also in agreement that myeloid lineage cells (*Itgam*^hi^ in non-B/T cells) were significantly increased in the state of obesity (*p* < 0.05), using the fraction of CD11b^+^ cells in CD45^+^ CD19^−^ CD3^−^ cells as a flow cytometry correlate (**Fig. 3i** and **3j**; detailed analysis for scRNA-seq data is available in **Code**).

### Spleen Cell Differences for Lean *versus* Obese Mice

We compared gene expression patterns in splenic immune cells between lean and obese mice. These cells have been shown to correlate with global immune cell dysregulation in clinical obesity^45,46^ ^47^ and, as a reservoir of blood cells, reflect immune cell types altered in other tissues due to obesity.^48,49^ Monocytes were the only cell type found to be altered in prevalence across the two conditions. This is in agreement with the observation of leukocytosis including increased blood monocytes in obese individuals.^45^ Differential expression (DE) in monocytes is shown in **Fig. 3f**, showing 50 genes with more than 2-fold change and *p* < 0.01 among 2011 genes identified for DE between the obese and lean groups (details available in **Code**). The most significantly upregulated gene in monocytes from obese mice was *Plin2* (**Fig. 3g**), which encodes a protein that forms a structural part of intracellular fat droplets and as such plays an important role in storage and handling of intracellular lipids. *Plin2* was reported to be upregulated in blood monocytes of obese humans.^50^ We also observed upregulation of *Pla2g7* which was reported as marker in atherosclerosis progression.^51^ For B and T cells, which did not shift in number between the lean and obese states, minimal DE differences were apparent across the conditions (**Fig. S13** and **S14**) with the exception of the upregulation of *Cpt1a* in B cells in obese mice. The *Cpt1a* gene product functions in mitochondrial beta-oxidation of fatty acids and its overexpression is consistent with previous reports of obesity.^47^

### Nanocoding of Heterogeneous, Debris-Rich Cells from Adipose Tissue (AT)

Digested AT is a particularly complex mixture often prepared as stromal vascular fraction (SVF) cells to deplete adipocytes. SVF contains diverse cell types, including immune cells, pre-adipocytes, fibroblasts, and endothelial cells, many of which are lipid-rich and buoyant, often with low viability (∼70%) after tissue digestion^52,53^ while ambient lipids and debris further complicate workup and analyses (**Fig. 4b** and **Fig. S15**).^53^ Investigations of SVF cell classes by multi-channel flow cytometry can be difficult due to a lack of surface markers, intense cell autofluorescence, cell-bound lipids, and lipid droplets.^54^ Current practices in SVF multiplexing for scRNA-seq rely on CD45 as a ‘pan-marker’^55^ which eliminates analysis of non-leukocyte cells that are critical for AT function (including preadipocytes, fibroblasts) as well as CD45^lo^ macrophage subtypes.^54,56^

**Fig. 4.**
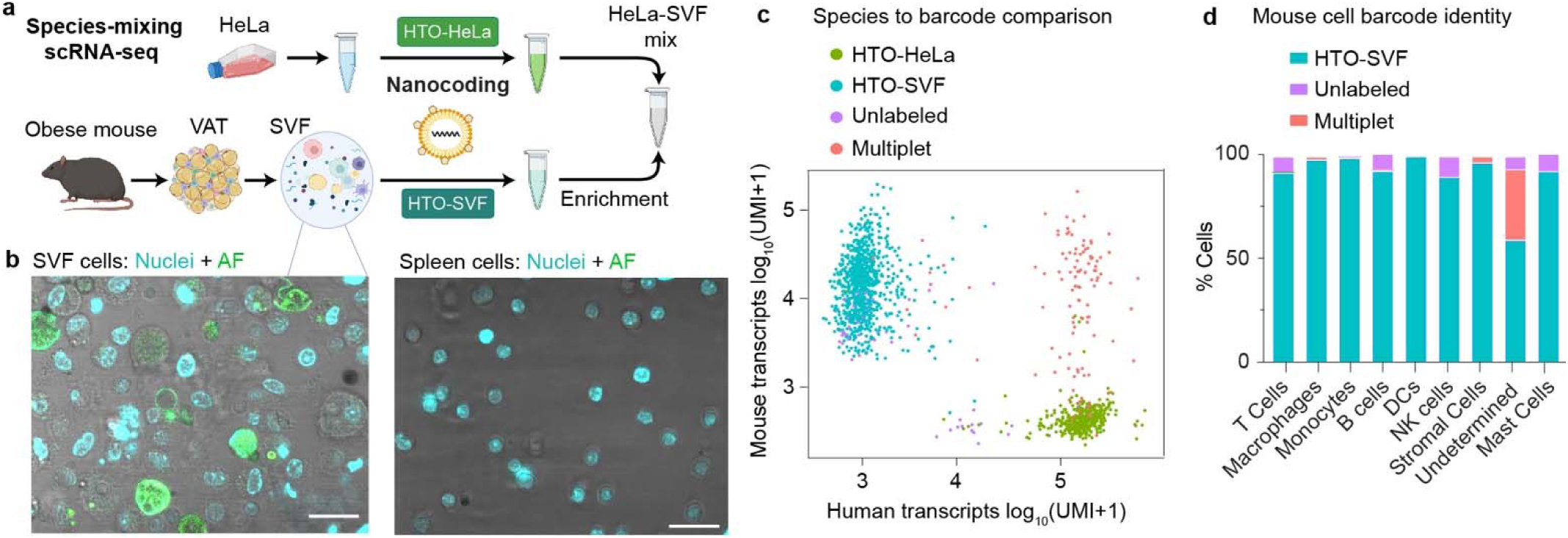
Nanocoding of heterogeneous mouse visceral adipose tissue (VAT) stromal vascular fraction (SVF). **(a)** ‘Species-mixing’ Nanocoding-based scRNA-seq experiment using a mixture of mouse SVF cells labeled with HTO-SVF barcode and human HeLa cells labeled with HTO-HeLa barcode before mixing. Barcodes are in **Table S1**. **(b)** Confocal fluorescence micrographs of SVF and spleen cells stained by nuclear dye (blue) with autofluorescence (AF) shown in green with overlay on brightfield image (grayscale). SVF contains more heterogeneous cells as well as small acellular structures, many of which are lipid-rich and cannot be fully eliminated. Scale bar: 20 μm. **(c)** Scatter plot showing human and mouse RNA counts clustered based on species origin with colors corresponding to identified barcodes determined with deMULTIplex. **(d)** Summary of labeling accuracy for mouse SVF cell types identified by the top variated genes from mouse RNA, distinguishing singlets (HTO-SVF) from multiplets and cells with undetected barcodes (unlabeled). Data processing steps are provided in **Code**. The marker genes to determine major immune cell types are listed in **Table S7**.

We used B-LNPs (3 μg/mL ssDNA) to label SVF cells, first using flow cytometry with fluorescent ssDNA which revealed strong labeling in both CD45^+^ and CD45^−^ populations (**Extended Fig. 1g**, **1h**, and **1i**). We note that B-LNPs based on ALC-0315 lipids which exhibited minimal biophysical labeling exhibited limited labeling of certain SVF cells, including ≥13% unlabeled CD45^+^ cells (**Fig. 1j** and **Fig. S16**). This emphasizes the advantage of multiple binding modalities for pan-cell labeling in heterogeneous samples. Using the MC3-based B-LNP, we then performed a two-plex Nanocoding ‘species-mixing’ experiment with mouse SVF and human cells as a ground truth for cell sample identification by species specificity of labels (**Fig. 4a**). The test revealed accurate barcode-based classification of human cells, mouse cells, and human-mouse multiplets (**Fig. S17**, **Fig. 4b**). However there was less bimodal separation in barcode count distributions for SVF cells compared with HeLa cells (**Fig. S17**). As a result, more SVF cells were negative for barcodes (4%, **Fig. 4c**) compared with cells from spleen (<0.5%). The unlabeled SVF cells were enriched for B, T, and NK cells (6–10% in **Fig. 4d**), potentially relating to the smaller sizes of lymphocytes.

Multiplexing was significantly improved through a 2-fold increase in B-LNP dose (6 μg/mL barcodes; **Fig. 5a**) which increased SVF barcode numbers ∼10-fold for minimal overlap between unlabeled and labeled cells (**Fig. 5b** and **Fig. S18**). We then used Nanocoding-based multiplexing to study AT cell gene expression changes related to aging using SVF cells from mice that were young (4 months, *N*=3) or aged (11 months, *N*=3). Both groups were on a high-fat diet which led to elevated body weight, especially in the aged mice (**Fig. S19**). We pooled the young and aged samples separately before Nanocoding to accommodate the extensive AT workup, despite the feasibility of 6-plex Nanocoding (*e.g*., **Fig. 3**). **Fig. 5c** shows a manual annotation of major cell types in all SVF based on top variated genes (**Fig. S20** and **Table S7**) using recent publications as references.^55,57–59,60,61^ After sample demultiplexing by barcode, more than 95% of cells were positively labeled in 11 of 12 manually annotated major cell types (**Fig. 5d**). Both CD45^−^ and CD45^lo^ cells exhibited high labeling (>95%; see **Fig. S21**), including fibroblasts, stromal cells, and epithelial cells, as well as some CD45^lo^ macrophages observed in recent publications.^56^ This pan-labeling capacity highlights the strengths of Nanocoding compared with more selective receptor-based barcoding with antibodies. Notably, only mast cells showed a high unlabeled population (29%) but these corresponded to just 8 of 5000 cells and ∼50% of these cells had low RNA quality (detected RNA sequence count less than 2000 per cell, **Fig. S22b**).

**Fig. 5.**
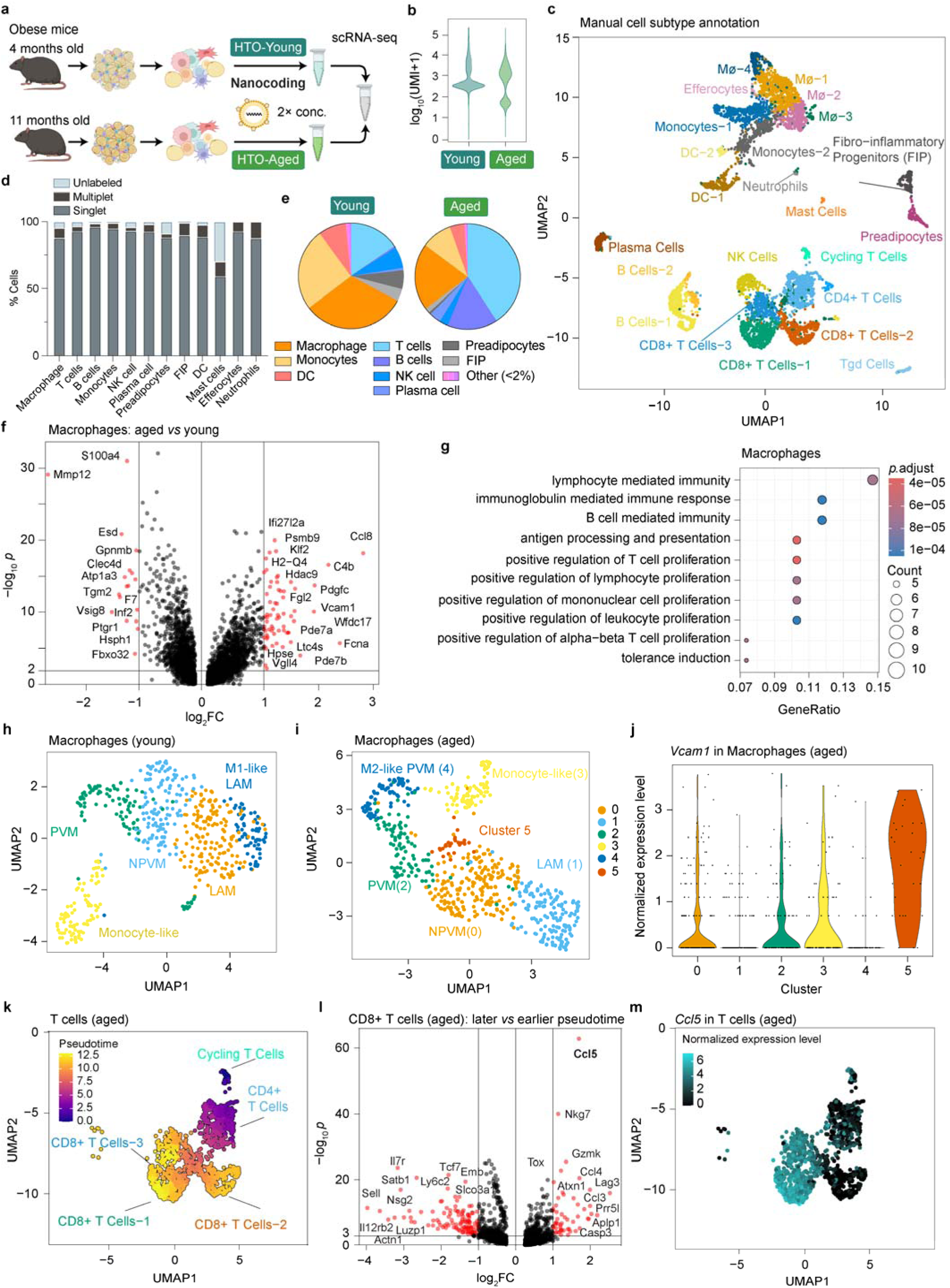
Application of Nanocoding for multiplexed scRNA-seq to analyze age-dependent gene expression in mouse VAT SVF cells. **(a)** Workflow for the young+aged adipose study, showing SVF cell collection from VAT of 4-month (‘young’) or 11-month (‘aged’) mice with diet-induced obesity using the same cell enrichment and sequencing workflow as in Fig. 4a but with twice the barcode concentration. **(b)** Raw counts per cell for the two barcodes, showing all cells identified in the experiment. The plot was generated with CellhashR. **(c)** UMAP of RNA-based cell clustering with manual annotation. The top variated genes for each cluster are shown in **Fig. S20** and marker genes are in **Table S7**. **(d)** Summary of labeling efficiency for each major cell type identified by RNA expression, distinguishing singlets and multiplets from those with undetected barcodes. Barcode classification was determined by deMULTIplex. **(e)** Proportions of major immune cell types in young and aged groups. **(f)** Differential gene expression of SVF macrophages, comparing aged *versus* young. Genes with more than 2-fold change (FC) and *p* < 0.01 are in red. Selected highly variated genes are labeled. **(g)** Ontology terms for genes upregulated in SVF macrophages in the aged group showing biological processes with the largest gene ratios. The dot size represents the number of genes with significant differential expression associated with the term, and the color represents the *p*-adjusted value. **(h,i)** UMAP of re-clustered macrophages from (**h**) young or (**i**) aged groups. LAM, lipid-associated macrophage; PVM, perivascular macrophage; NPVM, non-perivascular macrophage. The top variated genes for each cluster are shown in **Fig. S25** and **S26** and the list of marker genes is in **Table S9**. **(j)** *Vcam1* expression in each macrophage cluster from the aged group. The young group is shown in **Fig. S28**. **(k)** T cell pseudotime analysis in the aged group. Marker genes to determine each subtype are shown in **Fig. S31** and **Fig. S32**. **(l)** Differential gene expression of CD8+ T cells, comparing later pseudotime (clusters 1 and 3) *versus* earlier pseudotime (cluster 2). **(m)** *Ccl5* expression level overlay on the UMAP of T cells from the aged group. This cropped figure excludes several cells (**Fig. S33**).

### Differences between AT Cells from Young and Aged Obese Mice

Using Nanocoding-based scRNA-seq, we investigated age-related changes in AT SVF cells from obese mice. In the aged group, there was an increase in the fraction of lymphoid lineage cells (B and T cells) and a decrease in myeloid lineage cells (macrophages, DCs, neutrophils, and monocytes) (**Fig. 5e**), reflecting trends reported previously using flow cytometry.^62^ Gene expression differences in macrophages between aged and young mice are plotted in **Fig. 5f** adjacent to gene ontology representations in aged mice (**Fig. 5g**, **Fig. S23**; young data are shown in **Fig. S24**). In the aged group, a significant enhancement is observed in inflammatory-related molecular functions and biological processes which were not present in the young group, reflecting AT macrophage-driven inflammation in aged individuals. The macrophages were further independently clustered to explore subtypes, with manually annotated clusters shown in **Fig. 5h** and **5i** using markers from recent publications (**Table S9**).^55,57^ The fraction of perivascular macrophages (PVMs) and non-perivascular macrophages (NPVMs) were higher in the aged group, while lipid-associated macrophages (LAM) were lower in prevalence (**Fig. S27**). An additional cluster (cluster 5 in **Fig. 5i**) was observed only in the aged group with high expression of *Vcam1* (**Fig. 5j** and **Fig. S28**), a gene associated with inflammatory processes in atherosclerotic plaques,^63^ which may suggest a link between AT macrophages in aging with cardiovascular comorbidities of obesity. Additional analysis of preadipocytes (**Fig. S29**) and fibro-inflammatory progenitor cells (**Fig. S30**) are available in the **Supporting Information.**

Lymphoid lineage cells significantly increased in aged mice to become the dominant SVF cell population. A pseudo-time analysis of T cell clusters was performed in the aged group to evaluate their differentiation (**Fig. 5k**), with ‘early’ T cells designated as the ‘cycling T cell’ cluster classified by high expression of *Stmn1* and *Pclaf* (**Fig. S32**).^55^ Comparing CD8+ subclusters that are more differentiated (clusters 1 and 3) *versus* those less differentiated (cluster 2), the most significantly upregulated gene was *Ccl5* (FC > 3; (**Fig. 5l**), which encodes a potent T cell chemoattractant that could be related to the high T cell abundance in SVF from aged AT (**Fig. 5l**, **5m**, and **Fig. S33**). T cell exhaustion was specifically enhanced in the CD8+ T cell-1 cluster compared with other CD8+ clusters, reflected by high expression of *Lag3*, *Entpd1*, *Pdcd1, and Tigit* (**Fig. S34**). This high number of AT T cells is consistent with long-term inflammation in AT in chronic obesity with aging and their proposed function in resistance to treatments for sustained weight loss.^64,65^

## CONCLUSIONS

We conclude that Nanocoding using LNP-encapsulated ssDNA barcodes can label both cultured and primary cells efficiently and mildly without cell type dependency. The cell-specific labeling of these nanomaterials is enhanced compared with labels deriving from classical tools of chemical biology and molecular biology due to a combination of irreversible association processes driven by mechanisms controllable by the LNP composition and biochemical functionalization. Enhanced labeling is evident from both qPCR and multi-channel flow cytometry analyses. The stability of B-LNP can allow sample multiplexing with ample sample stability needed for processing after mixing cells with undetectable crosstalk. These features imply it is a robust pan-cell labeling method with minimal requirement in materials preparations or prior biological knowledge of samples to be tested, which are necessary as a multiplexing solution for broad sample types in scRNA-seq. The compatibility of Nanocoding with existing scRNA-seq platforms and the modular tunability of B-LNPs for cell labeling leaves substantial room for future labeling optimizations and precise selection of specific cell populations.

**Extended Fig. 1.**
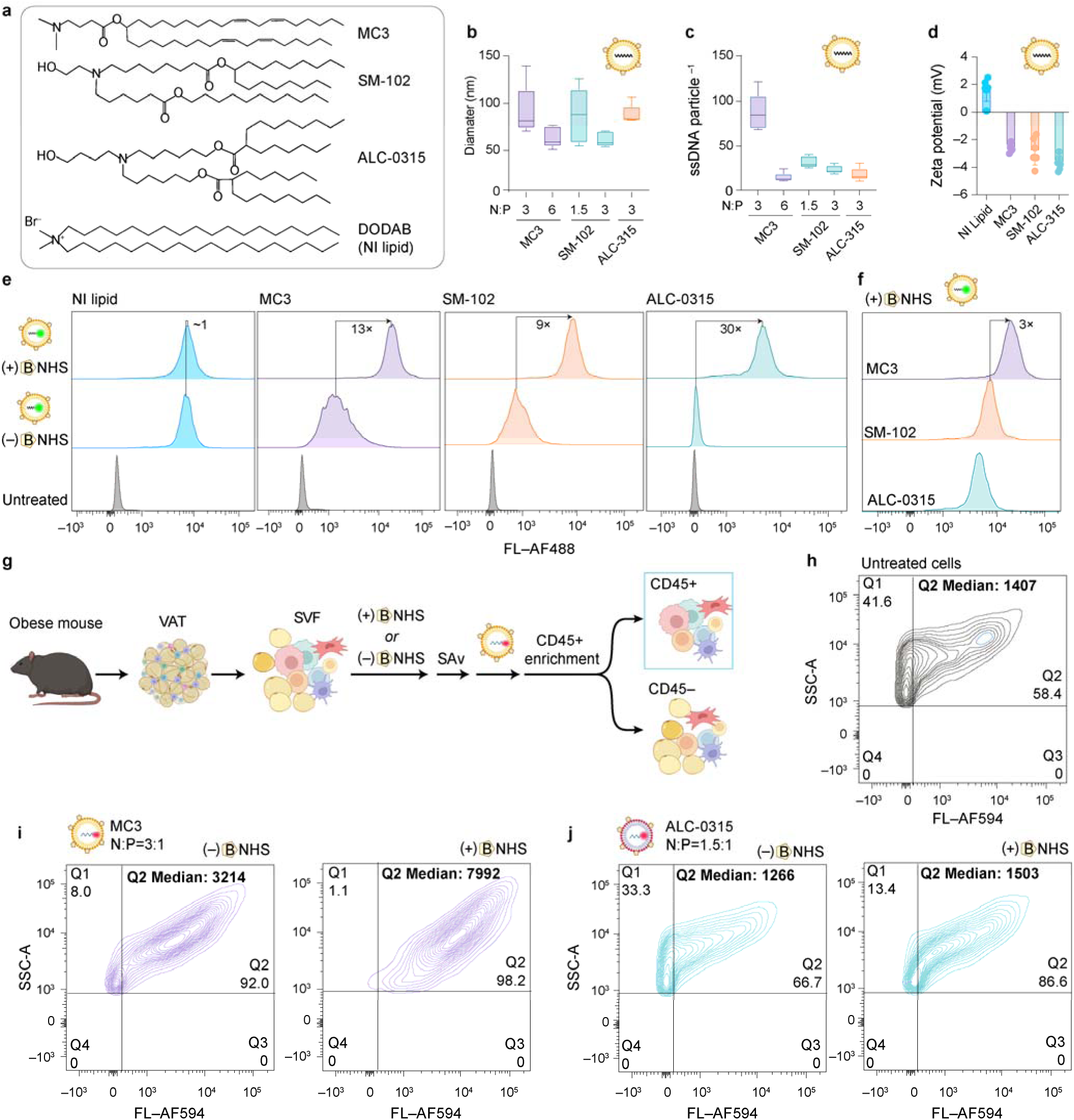
LNP composition control of physicochemical properties and cell labeling. **(a)** Chemical structures of three ionizable lipids tested, DLin-MC3-DMA (MC3), SM-102, and ALC-0315, and a non-ionizable (NI) lipid (DODAB). The other lipid components (B-PEG-DSPE, PEG-DMG, cholesterol, and DSPC) were included in fixed amounts (**Table S3**). **(b)** Hydrodynamic diameter of B-LNP with indicated composition of ionizable lipid, encapsulating AF488-DNA (**Table S1**), measured by FCS. **(c)** Mean number of ssDNA encapsulated per B-LNP particle using different ionizable lipids at indicated N:P ratios. **(d)** Zeta potential values for B-LNPs including the indicated lipid and N:P ratio. **(e)** HeLa cell labeling using the indicated B-LNP formulation at N:P = 3:1. **(f)** Cell labeling using B-LNP including MC3, SM-102, and ALC-0315 lipids, summarizing results from panel **e**. The fold change of median fluorescence of MC3-based B-LNP relative to that of ALC-012 is shown in the plot. **(g)** Schematic depiction of SVF cell collection and enrichment for CD45^+^ or CD45^−^ populations. Flow cytometry data are shown for CD45^+^ SVF cells, including **(h)** untreated cells to show autofluorescence, **(i)** cells treated with MC3-based B-LNP, and **(j)** cells treated with ALC-012-based B-LNP. Median AF594 intensity values of cells in Q2 are shown in the figure. Due to extensive autofluorescence of these cells, labeling is obscured but at least 13.4% of cells remain unlabeled with ALC-012-based B-LNPs, with media Q2 intensity similar to that of the autofluorescence background. The same gates were applied in each experimental group. Analysis of CD45^−^ populations is shown in **Fig. S16.**

## METHODS

### LNP encapsulation of ssDNA

To prepare ssDNA-encapsulated lipid nanoparticles, we adapted methods from previous reports.^28^ Stock lipids solutions in ethanol (30 or 90 µL) with total lipid concentration of 10 mg/mL were mixed at specified molar ratios listed in **Table S3** at a 1:3 volume ratio (ethanol: aqueous) with an aqueous solution (90 or 270 µL) containing ssDNA (∼80 μg/mL) in sodium citrate buffer (10 mM, pH 4) using a vortexer. To tune the ratio of ionizable lipid nitrogen-to-ssDNA phosphate (N:P ratio), the concentration of ssDNA and citrate buffer in the aqueous solution was adjusted while keeping all other volumes and concentrations fixed. The solution was then dialyzed against phosphate-buffered saline (PBS; pH 7.4) for 3 hours. The product was further purified by 10-fold concentration with a 100 kDa molecular weight cutoff (MWCO) centrifugal filter (Amicon Ultra; EMD Millipore, Billerica, MA) and then diluted 10-fold with PBS.^28^ This solution was stored at 4 °C until use.

### Cryogenic electron microscopy

To prepare samples for cryo-EM imaging, lacey carbon–coated 200 mesh copper grids (Electron Microscopy Sciences, PA) were glow discharged at 15 mA for 30 seconds with the PELCO easiGLOW Glow Discharge Cleaning System (Ted Pella, CA). Then, 3 μL of sample was applied to the grids. The grids were blotted with filter paper, and another 3 μL of sample was applied. The grids were then blotted and plunged into liquid ethane using a Vitrobot Mark IV System (Fischer, MA) at 4 °C and 90–95% humidity. The grids were kept in liquid nitrogen until imaging. Cryo-EM images are collected using EPU software on a Glacios Cryo-TEM (Thermo Fisher) at 200 kV with a Falcon4 Direct Electron Detector (Thermo Fischer).

### LNP size and zeta potential

Hydrodynamic size distribution was measured using dynamic light scattering (Malvern Panalytical, UK) and zeta potentiometry was conducted using folded capillary zeta cells (DTS1070, Malvern Panalytical).

### ssDNA encapsulation efficiency

The encapsulation efficiency (EE) of ssDNA in LNPs was determined by comparing the concentration of free ssDNA ([ssDNA]_free_) in the LNP-ssDNA solution relative to the total ssDNA including both free and encapsulated ([ssDNA]_total_) by releasing the ssDNA from LNPs with a 1:1 dilution with PBS containing 2% Tween-20 (Sigma Aldrich Inc., MO). The concentrations of ssDNA were determined *via* the Qubit ssDNA Assay Kit (Invitrogen, MA, USA) using a calibration curve was prepared using the barcode ssDNA as a standard in both PBS and PBS containing 1% Tween-20. After accounting for dilution factors, the encapsulation efficiency (EE) was calculated as

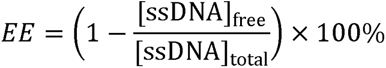

### Quantification of ssDNA copies per LNP

FCS was used to quantify dispersed LNPs that were fluorescently labeled after preparation with the dye DiO (0.1% in lipid mix; Invitrogen). The DiO-labeled LNPs were washed three times using 100 kDa MWCO centrifugal filters to minimize unbound DiO. LNP solutions were then diluted to ∼10 μg/mL by ssDNA concentration in PBS and added to 8-well glass-bottom LabTek chambers (Thermo Fischer). Fluorescence time traces were collected using an AlbaFCS instrument (ISS) with a diode laser (470 nm) for excitation and a single-photon avalanche photodiode detector for 10 s at a frequency of 100,000 Hz. The molar concentration of particles [LNP] was calculated by calibration with a standard fluorescent molecule (Rhodamine 110, Sigma-Aldrich) with known concentration, with model fitting applied based on our published protocols.^66^ The quantity of encapsulated ssDNA in each sample was determined as described above from the EE which allowed the calculation of the number of ssDNA per LNP as

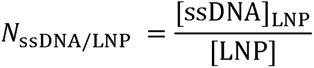

Where [ssDNA]_LNP_ = [ssDNA]_total_ – [ssDNA]_free_.

### Quantification of SAv binding to B-LNPs

A fluorescence-based assay was used to measure the SAv binding capacity of B-LNPs using AlexaFluor 647-labeled SAv (SAv-AF647; Invitrogen). First, LNPs (50 μL) or PBS (50 μL, control) were mixed with SAv-AF647 (0.2 mg/mL, 100 μL) in PBS containing 1% bovine serum albumin (BSA, R&D Systems, MN) for 15 minutes on ice. The mixture was then diluted to 1 mL with PBS and free SAv-AF647 was removed by filtration four times using a 100 kDa MWCO centrifugal filter. With each filtration cycle, the solution was centrifuged at 2000 × *g* for 8 minutes and the retentate (<200 μL) was diluted to 2 mL with PBS. For the final filtration step, PBS was added to the retentate to yield 200 μL of total volume and 100 μL was transferred to a flat-bottom 96-well black microplate (VWR International, PA, USA). The concentration of SAv-AF647 [SAv] was measured using a microplate reader (SpectraMax M5, Molecular Devices, CA), with calibration using a serial dilution of SAv-AF647 in the same plate. The number of SAv bound per B-LNP was determined by the following equation, using [LNP] determined above using FCS.

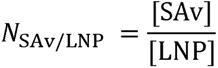

### Animals

All animal procedures were approved by the Institutional Animal Care and Use Committee (IACUC) at the University of Illinois Urbana-Champaign (protocol #22216). Four-week-old male C57BL/6J mice were purchased from The Jackson Laboratory (Bar Harbor, ME, USA). Upon arrival, mice were housed in groups of 4 in standard shoebox cages, individually ventilated in a temperature- and humidity-controlled environment, with 12-hour light:12-hour dark cycle. The diet-induced obesity (DIO) mice were fed a high-fat diet (60% kcal from fat [D12492]; Research Diets, Inc. New Brunswick, NJ) for a minimum of 12 weeks (young) or up to 11 months (aged). Lean control mice were fed a low-fat diet (10% kcal from fat [D12450J]; Research Diets, Inc.) for the same duration. Diet was refreshed twice weekly, and mice had access to fresh water and food *ad libitum*. Metabolic syndrome of DIO mice was confirmed by glucose tolerance test before tissue collection. At the appropriate timeline, mice (*N*=3/group) were sacrificed via CO_2_ asphyxiation and subsequent cervical dislocation, and tissues were collected for immediate downstream analyses.

### Cell isolation for six-sample spleen study

At the time of sacrifice, DIO mice and age-matched lean mice were 8 months old. Freshly collected spleens were minced with scissors and passed through a 70 μm filter (Corning, NY) with a rubber plunger before cells were collected as the filtrate in PBS. The cells were treated with RBC Lysis Buffer (eBioscience, Invitrogen) for 5 min on ice and then washed twice with HBSS buffer (Corning, NY) containing 2% BSA and centrifuged at 500 × *g* for 5 min at 4 °C. The supernatant was discarded, and cells were resuspended in HBSS (10 mM without Ca^2+^, Corning, NY) for Nanocoding and flow cytometry steps on the same day.

### Cell isolation for young+aged adipose study

At the time of sacrifice, aged obese mice were 11 months old and young obese mice were 4 months old. Fresh gonadal and perirenal adipose tissues were minced with scissors and incubated in collagenase type II (2 mg/mL; StemCell Technologies, CA) and DNase I (20 µg/mL; Sigma-Aldrich Inc.) in HBSS buffer (10 mM with Ca^2+^). Tissue was incubated at 37 °C at 80 r.p.m. for 45 minutes. When tissue pieces were no longer visible, the suspension was passed through a 100 µm cell strainer and centrifuged at 500 × *g* for 10 minutes at 4 °C. Floating adipocytes and supernatant were removed and the cells were treated with RBC Lysis Buffer for 5 min on ice. Then cells were washed twice with HBSS buffer containing 2% BSA and centrifuged at 500 × g for 5 min at 4 °C. The supernatant was discarded and cells were resuspended in HBSS (10 mM without Ca^2+^) for Nanocoding steps on the same day.

### Cell Culture

HeLa or RAW 264.7 cells were cultured in Dulbecco’s Modified Eagle Medium (Corning, NY) supplemented with 10% fetal bovine serum (Gibco, MA) and 0.01% penicillin/streptomycin (Thermo Fisher) at 37 °C in 5% CO_2_. Cells were harvested from adherent cultures using TrypLE Express (without Phenol Red, Gibco, Thermo Fisher) and suspended in HBSS (10 mM with Ca^2+^, Corning) for Nanocoding steps on the same day.

### Nanocoding with B-LNPs

Cells were suspended at 10^6^ to 10^7^ per milliliter in PBS or HBSS (without BSA) and mixed with NHS-PEG_4_-biotin (B-NHS; Cayman Chemical, MI) at 100–200 μM for 15 minutes on ice. The cells were then centrifuged (350–400 × g, 4 °C, 4 minutes) and washed with PBS and optionally PBS containing 1% BSA. After washing, the cells were resuspended in PBS with 1% BSA and mixed with SAv (2–3 g/mL). The cells were incubated for 5 minutes on ice and then centrifuged to remove the SAv-containing supernatant. Then, a solution of B-LNP (∼4 μg/mL by barcode ssDNA) was added at a 1:1 volume ratio to the cell suspension. The mixture was incubated for 15 minutes on ice and the cells were washed twice by centrifugation (350-400 × *g*, 4 °C, 4 minutes) using PBS for resuspension. The final cell suspension in PBS was near 10^6^ mL^−1^. At this stage, spleen cells and cultured cells were ready for scRNA-seq. For SVF cells, we performed live cell enrichment using a Dead Cell Removal kit (Miltenyi Biotec, Germany) followed by CD45+ enrichment using Mouse CD45 Microbeads (Miltenyi Biotec). This CD45-selective enrichment process was performed instead of sorting so as to not fully exclude CD45^−^ and CD45*^lo^* populations to benchmark labeling of diverse cell types depleted by sorting.^55–57^

### Flow Cytometry

For fresh spleen cells, after RBC lysis and wash steps, cells were suspended in PBS with 1% BSA and stained with fluorophore-conjugated anti-mouse antibodies and viability dyes for 25 minutes at 4 °C (full list and dilution factors in **Table S5**). An anti-CD45 antibody (BV605 conjugate), anti-CD3 antibody (PE conjugate), anti-CD19 antibody (FITC conjugate), and anti-CD11b antibody (AlexaFluor647 conjugate) cocktail was used for the spleen-cell-mixing test shown in **Figure 3**. RAW 264.7 cells were stained with antibodies using the same procedure, or stained with DiO (1 mM in DMSO, 1:200 dilution, Invitrogen) for 15 minutes at room temperature. Cells were washed twice with PBS containing 1% BSA and analyzed using a BD Symphony A1 (BD Biosciences, NJ). Data were analyzed with FlowJo software (version 10.9.0).

### qPCR

To evaluate barcode labeling of cells, total DNA of cultured cells or spleen cells was isolated using the DNeasy Blood & Tissue Kit (Qiagen, Valencia, CA, USA). A spike-in sequence was added prior to DNA extraction at the same concentration in each sample. DNA concentrations were determined using an ND-1000 spectrophotometer (Nanodrop Technologies, Wilmington, DE, USA). Each sample was diluted 10-fold and 3 μL was used for each qPCR test. Each sample was mixed with primers (1.5 μL of each primer per test) and PCR mastermix containing SYBR reagent (5 μL per test; Applied Biosystems Power SYBR Green PCR Master Mix 2× Fisher Scientific). A total of 10 μL was used for each test. All gene expression data were analyzed on a Thermo Fisher Connect Platform using the ΔΔC_T_ method, with barcode sequence levels calculated relative to the spike-in sequence.

### scRNA-seq

#### Construction of 10× 3’GEX+CSP libraries

Single-cell libraries were prepared at the DNA Services laboratory of the Roy J. Carver Biotechnology Center at the University of Illinois Urbana-Champaign. Ten thousand single-cell suspensions showed an average viability of 90% for the six-sample spleen study and 86% for the young+aged adipose study by acridine orange/propidium iodide staining on the Nexcelom K2 (Nexcelom Bioscience, MA). The suspensions were converted into a barcoded gene expression library (GEX) and a barcoded cell surface protein (CSP) library with the Chromium Next GEM Single-Cell 3’ Dual-Index Kit (v3.1) with Feature Barcode technology for Cell Surface Protein (10X Genomics, CA) following the manufacturer’s protocols.

The 10X Chromium instrument separates thousands of single cells into Gel Bead Emulsions (GEMs) that add a barcode to the mRNA and cell surface protein from each cell. Following ds-cDNA synthesis and DNA amplification from cell surface protein Featured Barcodes, a barcoded, individual 3’ GEX library and CSP library compatible with the Illumina chemistry were constructed. The final libraries were quantitated by Qubit (Thermo Fisher) and the average size was determined on the AATI Fragment Analyzer (Agilent, CA). The libraries were diluted to 5 nM concentration and further quantitated by qPCR on a Bio-Rad CFX Connect Real-Time System (Bio-Rad, CA).

#### Sequencing

The final 10X single cell GEX library and CSP library were sequenced on one lane in an Illumina NovaSeq X Plus 10B Flowcell (Illumina, CA) as paired reads with 28 cycles for read 1, 10 cycles for each index read, and 150 cycles for read 2. Data in resulting fastq.gz files were demultiplexed with bcl-convert v 4.2.4 software from Illumina.

### Bioinformatic Analysis

The sequenced RNA and barcode libraries were aligned and counted using CellRanger 7.1.0 based on references.^67^ The ‘species-mixing’ test used a combined reference of human and mouse genes (mouse: GRCm39, human: GRCh38). The resulting count matrices were processed with Seurat v5.0.3.^68^ Cells were then filtered based on percent mitochondrial value as 3 times the median absolute deviation (MAD) away from the median, using the isOutlier function from the Scuttle v1.12.0 package.^69^ For the six-sample spleen study, the data matrix after CellRanger count was normalized and clustered using the Seurat v5.0.3 and loaded for the downstream gene analysis. For samples containing SVF cells, the data matrix was additionally run through SoupX to remove ambient RNA before a second cycle of normalization and clustering, following previous practices for SVF samples.^18,55^

Cell clusters were manually annotated based on the expression of known marker genes (reference database: PanglaoDB) for SVF cell sample, or automatically annotated by SingleR v2.4.1 using ImmGen or MouseRNAseq as databases for the six-sample spleen study.^43,44,59,70^ The HTO barcode count matrix was filtered to match the filtered RNA count matrix, and then run through CellhashR v1.1.0 for raw barcode count quality check and sample classification.^19^ The barcode matrix was combined with the RNA matrix as a single Seurat object for the following analysis.

The differential gene expression analysis was run using default methods in Seurat or pseudo-bulking methods.^71^ Genes with selected significance were evaluated through Gene Ontology (GO) using clusterProfiler v 4.10.1.^72,73^ For macrophages in the young+aged adipose study, they were subset from the original dataset and separated by HTO-Aged or HTO-Young labels to run clustering independently. For T cells, the manually identified T cell subgroups were subset as single Seurat objects and loaded to Monocle3 v 1.3.7 for pseudotime analysis.^74,75^ Detailed steps of the analysis are available in **Code**.

## Acknowledgments

This work was supported by funds from the National Institutes of Health (R01DK112251, R01DK124290, R01DK127531, R01DK131782, R35HL167143). C.C.A. was supported by the Beckman Foundation. N.Y.G.M. was supported by a fellowship from the National Science Foundation. O.H.A. acknowledges support from the National Institutes of Health under award number T32EB019944. We thank the DNA service group at Roy J. Carver Biotechnology Center at the University of Illinois Urbana-Champaign for help in library preparations and sequencing. We thank the High-Performance Biological Computing Center (HPCBio) at the University of Illinois Urbana-Champaign for suggestions in bioinformatic processing steps. We thank the Materials Research Laboratory (MRL) at the University of Illinois Urbana-Champaign for help in cryo-EM data. Fig. 1, Fig. 3a, Fig. 4a, Fig. 5a, and schematic symbols for LNPs or antibodies were created using components from BioRender.com.

## Author Contributions

Y.F. and A.M.S conceptualized the work, designed experiments, and developed the protocols. Y.F. did LNP formulation, LNP characterizations, analytical flow cytometry, and qPCR experiments. Y.F. and D.C. did tissue digestion experiments and optimizations. C.C.A. and N.Y.G.M provided animal tissues. Y.F., C.L.W., and F.X. performed scRNA-seq experiments. Y.F. and J.D. performed bioinformatic analysis. O.H.A. and Y.F. performed microscopy tests. C.K. and Y.F. performed FCS tests. Y.F. and A.M.S. wrote the manuscript.

## Conflict of Interest

The authors declare no competing financial interests.

## Data

Raw and processed sequencing data generated in this study have been submitted to GEO (accession number: GSE280262). The human reference genome (GRCh38) and the mouse reference genome (GRCm39) were from https://useast.ensembl.org/index.html.

## Code

Scripts and code are available via GitHub at https://github.com/yujunf2/Spleen-6-sample-multiplex.git (Spleen-sample), https://github.com/yujunf2/HeLa-SVFmix-scseq.git (HeLa-SVF-sample), and https://github.com/yujunf2/SVF-sample-Young-Aged-mixed.git (SVF cell Young-Aged Sample). Scripts and code are available via GitHub at https://github.com/yujunf2/Spleen-6-sample-multiplex.git (Spleen-sample), https://github.com/yujunf2/HeLa-SVFmix-scseq.git (HeLa-SVF-sample), and https://github.com/yujunf2/SVF-sample-Young-Aged-mixed.git (SVF cell Young-Aged Sample).

## Notes

https://www.ncbi.nlm.nih.gov/geo/query/acc.cgi?acc=GSE280262

### Competing Interest Statement

The authors have declared no competing interest.

