## Supporting Information for "Nanocoding: Lipid Nanoparticle Barcoding for Multiplexed Single-Cell RNA Sequencing"

#### Table of Contents

1 Supporting Figures …………………………………………………………… 3

2 Supporting Tables ……………………………………………………………. 37

3 Supporting References ……...………………………………………..…... 47

### 1 Supporting Figures

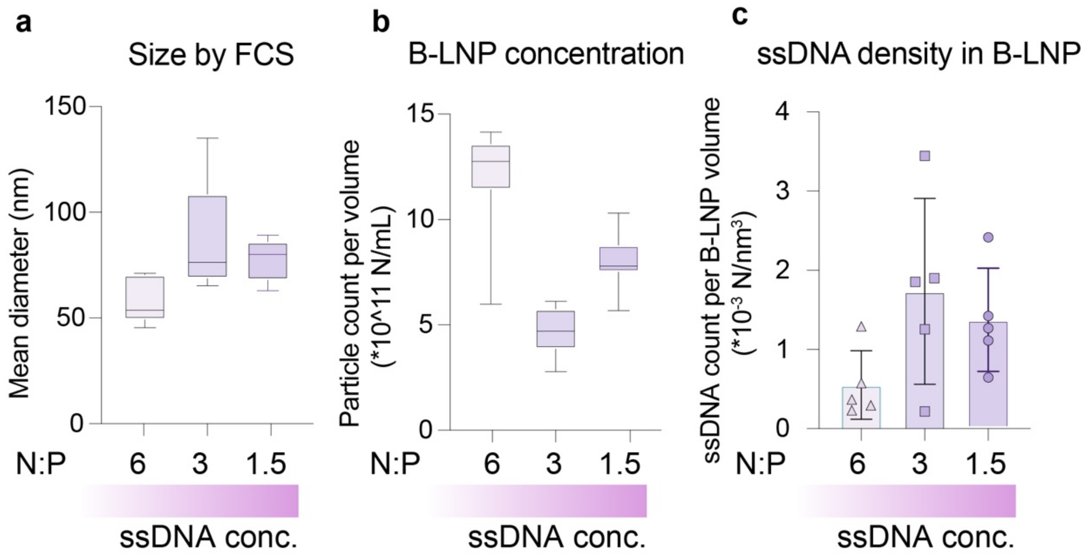

**Figure S1.** Size and ssDNA encapsulation of B-LNPs at different ratios of lipid nitrogen to DNA phosphate (N:P) using LNPs containing MC3-based on fluorescence correlation spectroscopy (FCS) measurements. **(a)** Size of B-LNP by FCS. **(b)** B-LNPs particle number per solution volume by FCS. **(c)** Mean ssDNA number per B-LNP volume.

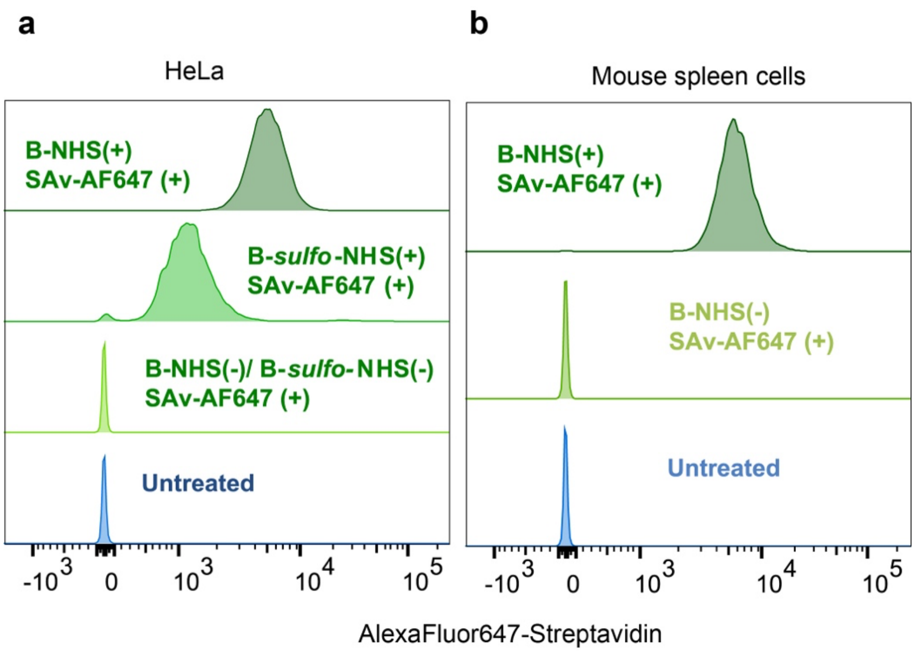

**Figure S2.** Characterization of NHS-based reagents for cell surface biotinylation using Streptavidin-AlexaFluor647 (SAv-AF647) conjugates measured by flow cytometry. **(a)** Labeling of HeLa cells using *sulfo*-NHS-biotin (B-*sulfo*-NHS) with 4-carbon linker between biotin in comparison with NHS-PEG$_4$-biotin (B-NHS) which has a tetraethylene glycol spacer. Labeling is enhanced 4-fold with B-NHS. SAv-AF647 treatment without surface biotinylation yields similar signals compared with untreated cells. **(b)** B-NHS labeling of cells from spleen digests. SAv-AF647 treatment without surface biotinylation yields similar signals compared with untreated cells.

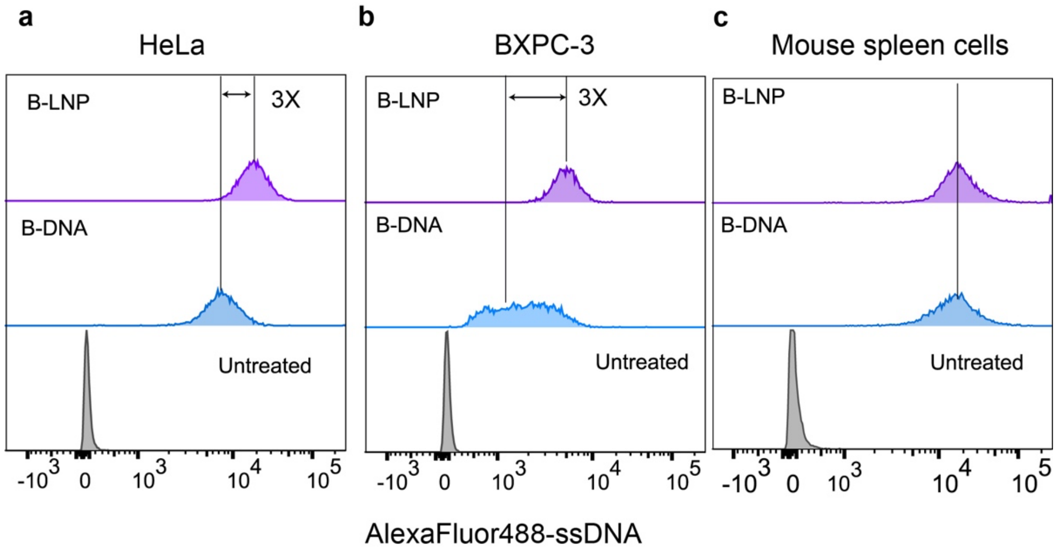

**Figure S3.** Cell labeling comparison between biotin-ssDNA conjugate (B-DNA) and B-LNP. Flow cytometry was used to measure AF488-DNA-Biotin (**Table S1**) for B-DNA labeling and AF488-DNA (**Table S1** and **S3**) for B-LNP labeling. The AF488-DNA concentration was 3 µg/mL for both label classes. In phosphate-buffered saline, the quantum yield of B-LNP-AF488 was 33% relative to that of B-DNA-AF488, so B-LNP-based labeling could be underestimated by up to 3-fold based on fluorescence intensity. Cells include **(a)** HeLa, **(b)** BXPC-3, and **(c)** mouse spleen cells. Fold change of median fluorescence intensity is indicated in each graph.

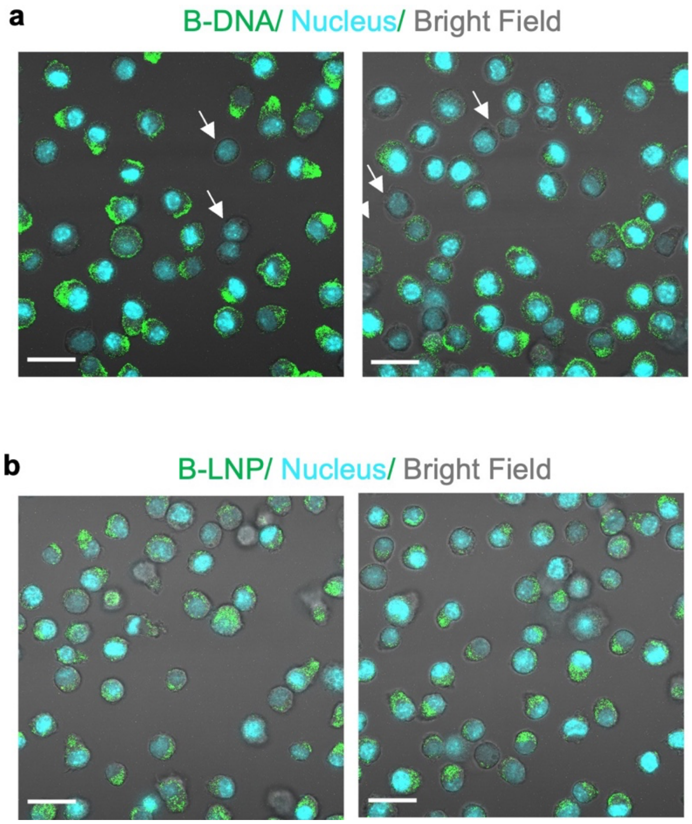

**Figure S4.** Cell labeling comparison between B-DNA and B-LNP by confocal microscopy. RAW 264.7 cells were labeled with **(a)** B-DNA-AF488 or **(b)** B-LNP containing AF488-DNA and imaged by confocal microscopy. Two images from the same sample are shown for each group. The AF488 fluorescence channel (green) and nucleus stain (blue) are overlayed on the brightfield image (gray) to show the difference in B-DNA or B-LNP label distribution in cells. The arrow indicates cells that are in focus but apparently unlabeled by B-DNA. Scale bar: 20 μm.

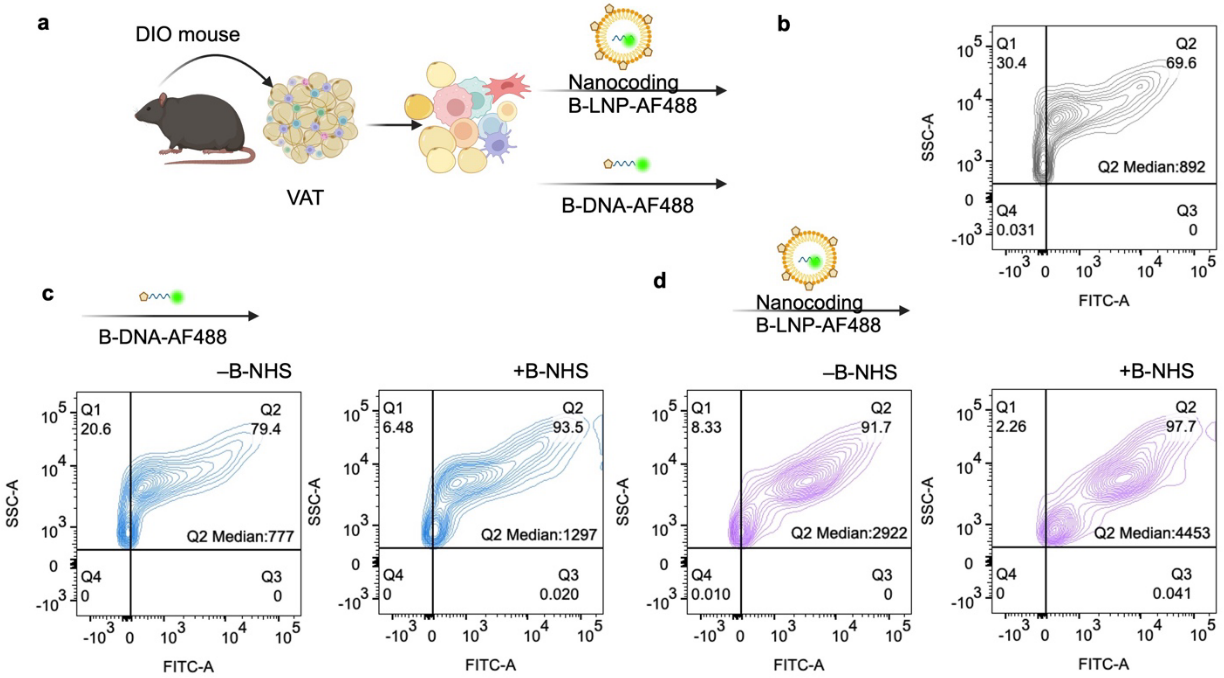

**Figure S5.** Comparison between B-DNA and B-LNP for labeling heterogeneous mouse visceral adipose tissue (VAT) stromal vascular fraction (SVF). **(a)** Schematic depiction of SVF cell collection and labeling with B-LNP or B-DNA. Cells were first labeled with B-NHS and SAv (not shown) before the addition of B-LNP containing AF488-DNA or B-DNA-AF488 with equal amounts of ssDNA (3 µg/mL). Flow cytometry data are shown for SVF cells, including **(b)** untreated cells to show autofluorescence, **(c)** cells treated with B-DNA, and **(d)** cells treated with B-LNP. The median AF488 intensity value of cells in Q2 is shown in each figure panel. Due to the extensive autofluorescence of these cells, labeling is obscured but at least 6.5% of cells remain unlabeled with B-DNA, with mean intensity similar to that of the autofluorescence background. The median intensity of cells in Q2 is higher for B-LNP (4453) *versus* B-DNA (1297). The quantum yield of B-LNP is 33% relative to B-DNA in phosphate-buffered saline so B-LNP-based labeling could be underestimated by up to 3-fold based on fluorescence intensity.

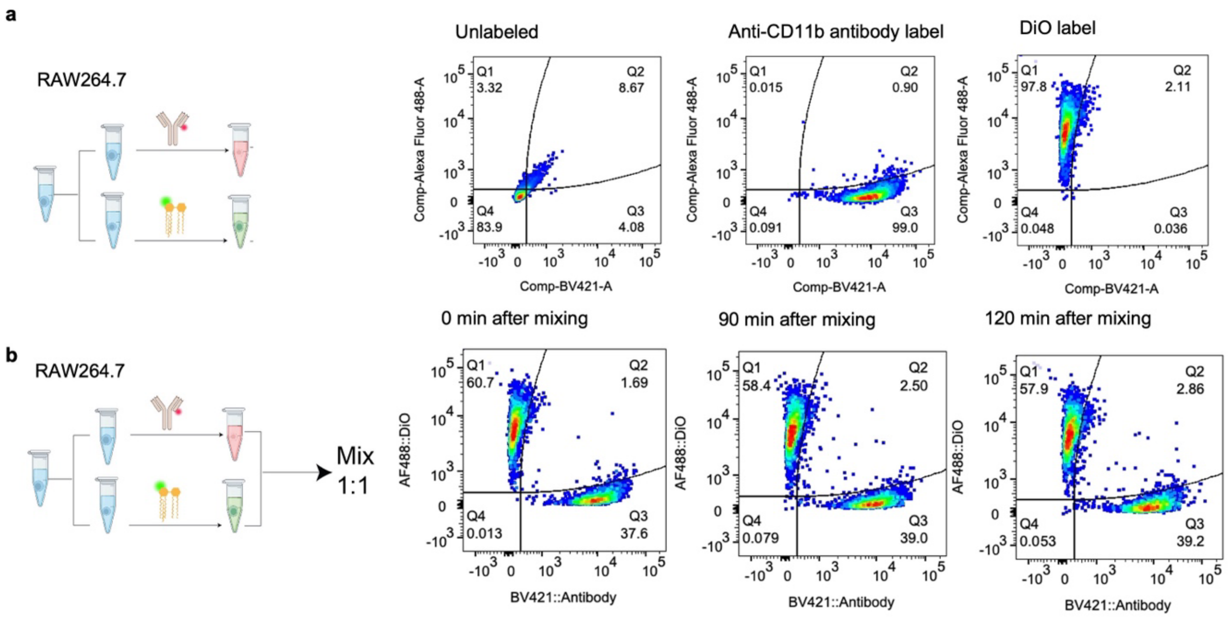

**Figure S6.** Labeling stability test using RAW 264.7 cells and multi-channel flow cytometry. Cells were labeled with an anti-CD11b primary antibody (BV421 conjugate) or DiO (**Table S5**), then mixed and tested for changes in fluorescence intensity over time based on the percentage of cells within four gates (Q1–Q4), shown in each plot. Dye transfer is measured by the cell population in the Q2 gate. In particular, cells near the Q1/Q2 interface or the Q2/Q3 interface, respectively, represent the transfer of antibody or DiO label. **(a)** Fluorescence intensities of untreated cells and those labeled with only the antibody or DiO labels. **(b)** Fluorescence intensities after mixing 1:1 for the indicated times. Samples were kept on ice after mixing.

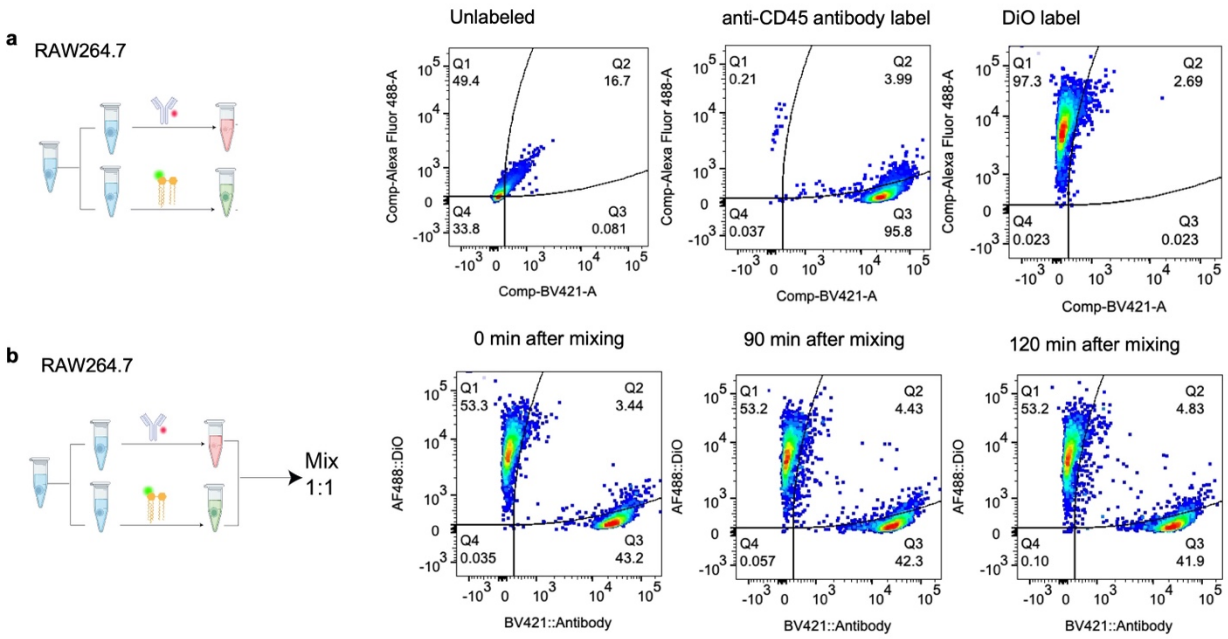

**Figure S7.** Cells were labeled with an anti-CD45 primary antibody (BV421 conjugate) or DiO (**Table S5**), then mixed and tested for changes in fluorescence intensity over time based on the percentage of cells within four gates (Q1–Q4), shown in each plot. Dye transfer is measured by the cell population in the Q2 gate. In particular, cells near the Q1/Q2 interface or the Q2/Q3 interface, respectively, represent the transfer of antibody or DiO label. **(a)** Fluorescence intensities of untreated cells and those labeled with only the antibody or DiO labels. **(b)** Fluorescence intensities after mixing 1:1 for the indicated times. Samples were kept on ice after mixing.

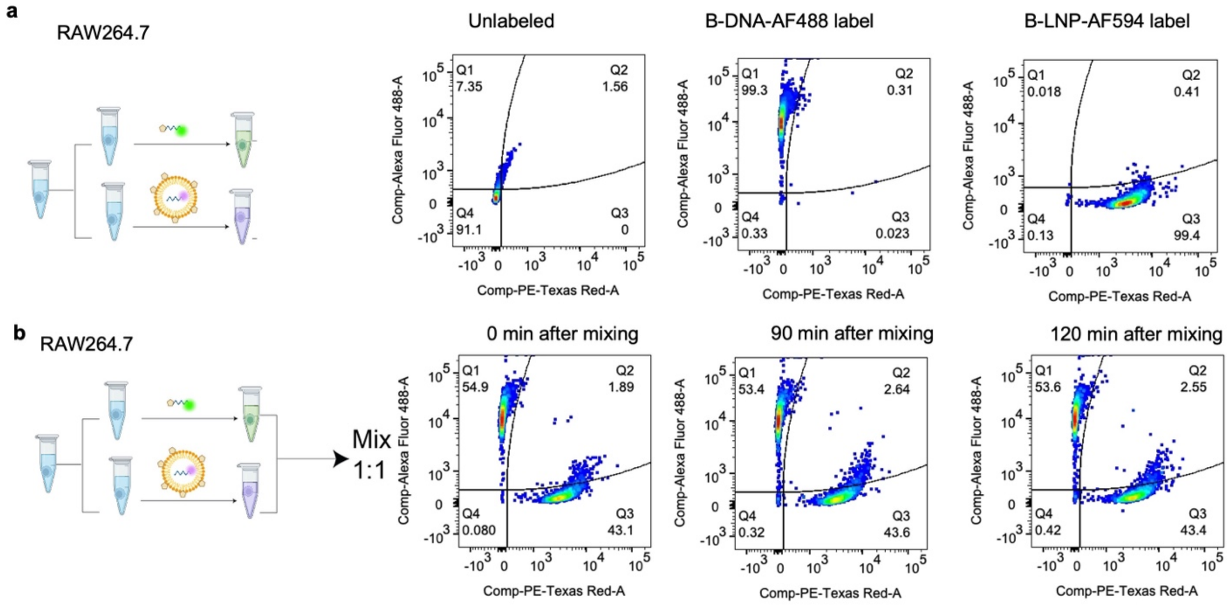

**Figure S8.** Labeling stability test using RAW 264.7 cells and multi-channel flow cytometry. Cells were labeled with B-LNP (encapsulating AF594-DNA) or B-DNA (AF488-ssDNA-Biotin) (**Table S1**), then mixed and tested for changes in fluorescence intensity over time based on the percentage of cells within four gates (Q1–Q4), shown in each plot. Dye transfer is measured by the cell population in the Q2 gate. In particular, cells near the Q1/Q2 interface or the Q2/Q3 interface, respectively, represent the transfer of B-LNP or B-DNA label. **(a)** Fluorescence intensities of untreated cells and those labeled with only B-LNP or B-DNA. **(b)** Fluorescence intensities after mixing 1:1 at the indicated times. Samples were kept on ice after mixing.

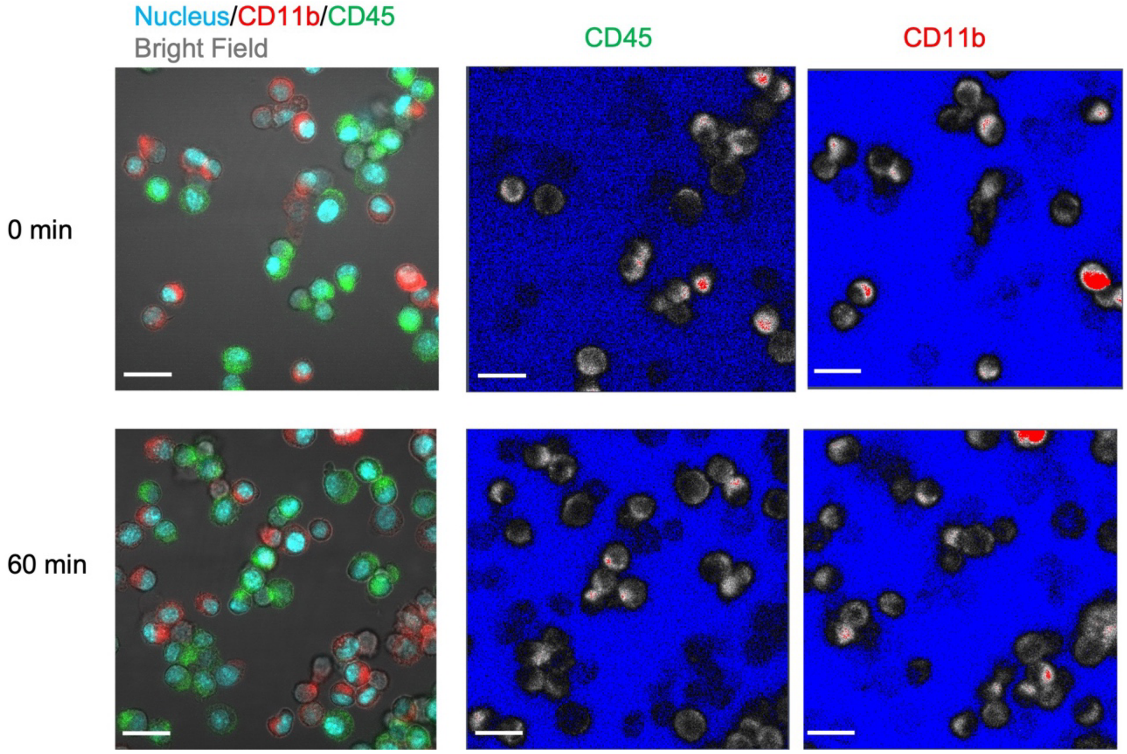

**Figure S9.** Antibody-based labeling stability test using RAW 264.7 cells and fluorescence microscopy. Cells were labeled with anti-CD11b antibody (AF647 conjugate) or anti-CD45 antibody (AF488 conjugate) (**Table S5**), then mixed 1:1. Cells were imaged by confocal microscopy over time. The left images show an overlay of anti-CD11b (red), anti-CD45 (green), nuclear stain (blue), and brightfield (gray). The images on the right show anti-CD11b and anti-CD45 channels alone in Range Indicator mode (minimum in blue and maximum in red) in Zen Lite software. The anti-CD45 label can be observed to transfer to anti-CD11b cells after 60 minutes compared with the initial conditions (“0 min” was imaged within 10 minutes of mixing). Scale bar: 20 μm.

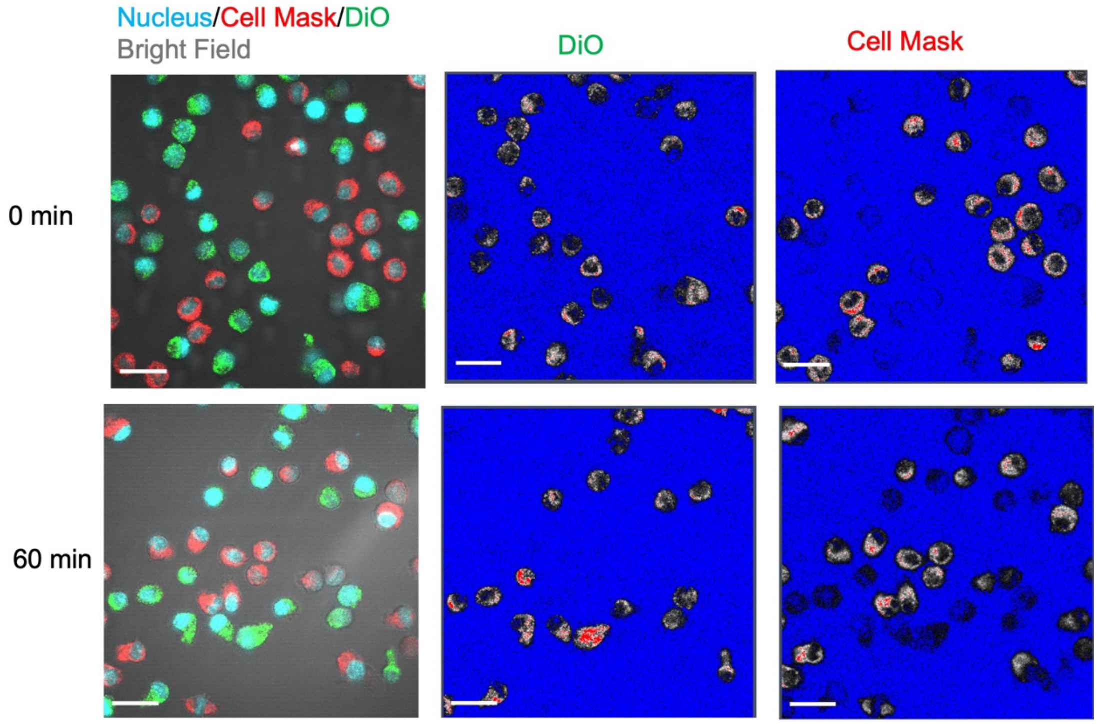

**Figure S10.** Lipid-based labeling stability test using RAW 264.7 cells and fluorescence microscopy. Cells were labeled with one of two lipid-based dyes with distinguishable colors (Cell Mask Orange or DiO) (**Table S5**), then mixed 1:1. Cells were imaged by confocal microscopy over time. The left images show an overlay of Cell Mask (red), DiO (green), nuclear stain (blue), and brightfield (gray). The images on the right show Cell Mask and DiO channels alone using Range Indicator mode (minimum in blue and maximum in red) in Zen Lite software. Cell Mask label can be observed to transfer to DiO-labeled cells after 60 minutes compared with the initial conditions (“0 min” was imaged within 10 minutes of mixing). Scale bar: 20 μm.

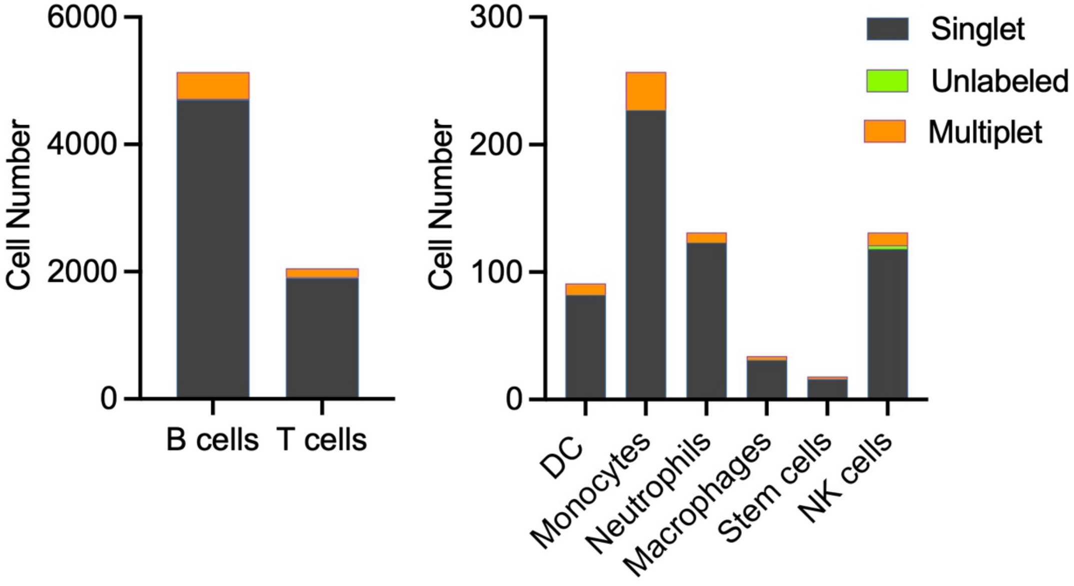

**Figure S11.** Barcode identification in 8 cell types from six-sample spleen study. Plots distinguish cells positively identified for a barcode as a single or multiplet as well as those remaining unlabeled.

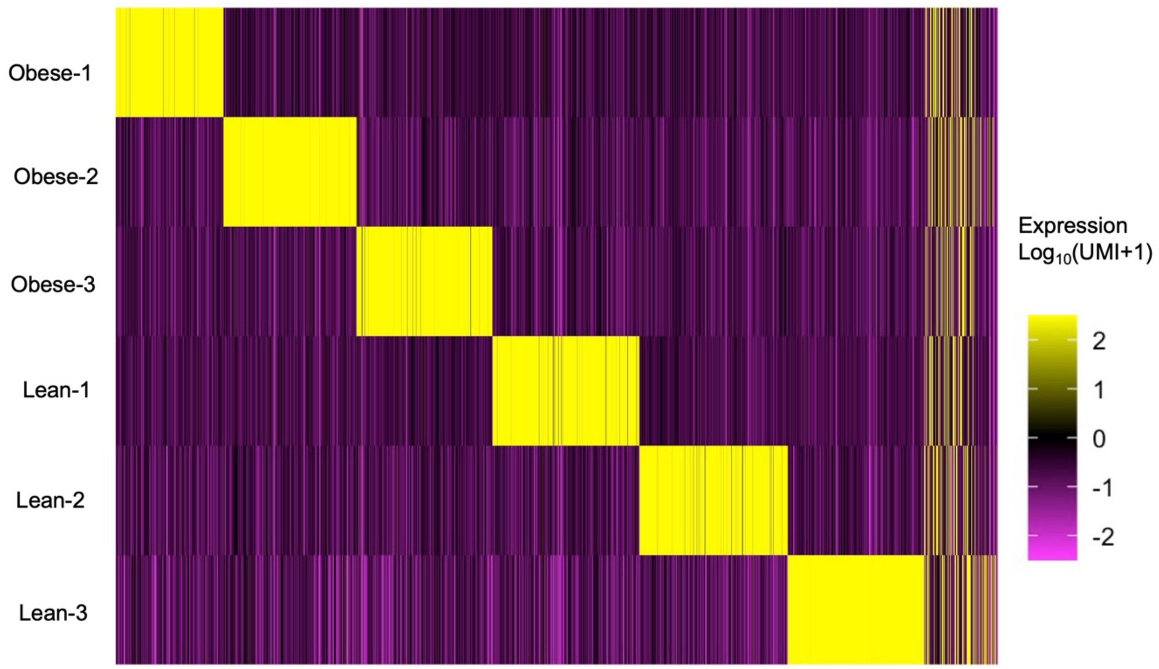

**Figure S12.** Heatmap representation of barcode counts in six-sample spleen study using deMULTIplex. Reads are shown for barcode sequences (6 rows) in 8019 cells (thin columns, after cell quality filtering) sorted by deMULTIplex and plotted using the CellhashR package.[1,2]

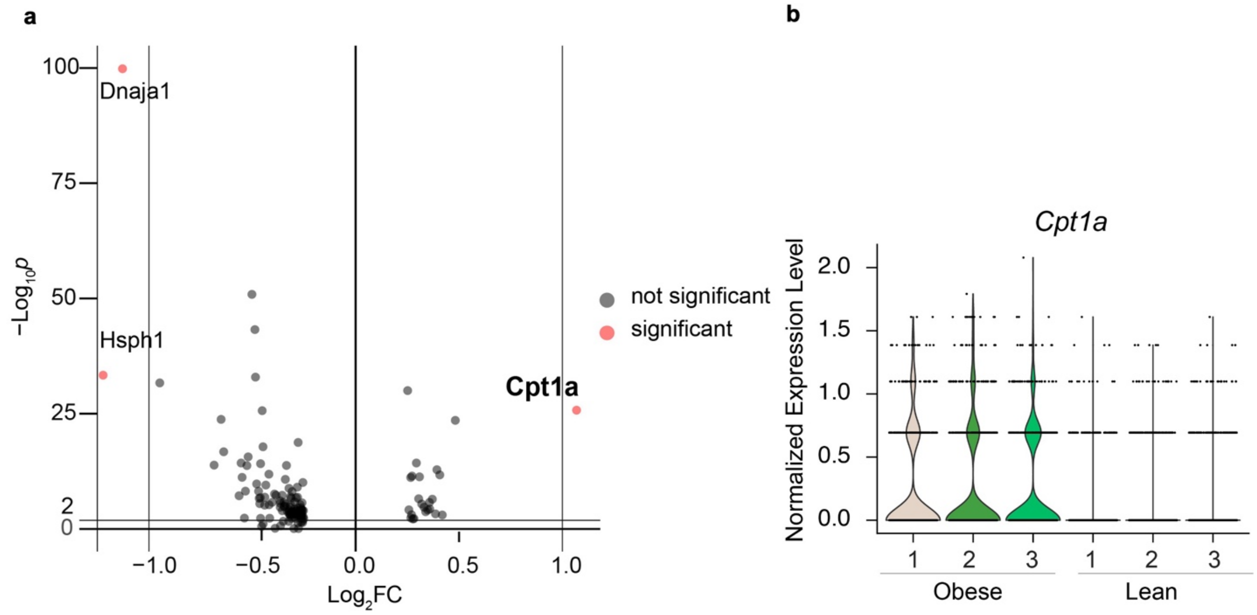

**Figure S13.** Differential gene expression from six-sample spleen study showing B cells from spleens of lean and obese mice. **(a)** Differential gene expression in B cells comparing obese *versus* lean mice, aggregating all samples in each biological condition. Selected top variated genes are labeled. Those in bold text have $p < 0.001$ by pseudobulk analysis. **(b)** *Cpt1a* expression in each sample with gene counts normalized and variance stabilized by SCTransform in the Seurat v5 package and pseudobulk analysis conducted using DESeq2.[3,4]

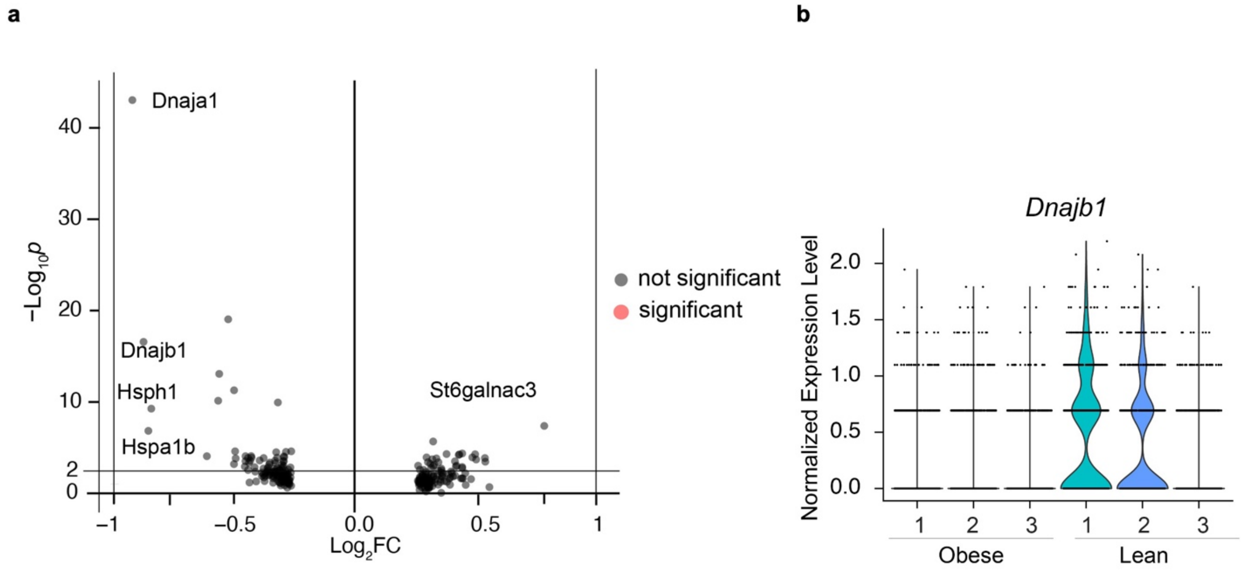

**Figure S14.** Differential gene expression from six-sample spleen study showing T cells from spleens of lean and obese mice. **(a)** Differential gene expression of T cells comparing obese *versus* lean mice, aggregating all samples with each biological condition. Selected top variated genes are labeled. Those in bold text have $p < 0.001$ by pseudobulk analysis. **(b)** *Dnajb1* expression in each sample with gene counts normalized and variance stabilized using the Seurat v5 package and pseudobulk analysis conducted using DESeq2.[3,4]

Mouse tissue digest

Adipose Spleen

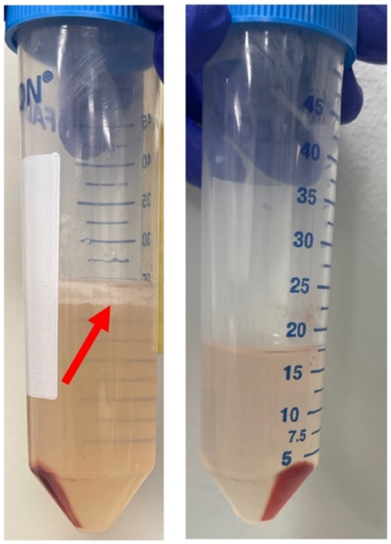

**Figure S15.** Difference between digested adipose tissue and spleen tissue after centrifugation. The red arrow indicates buoyant lipids in the adipose tissue digest. The red pellets in each sample are primary cells before red blood cell lysis.

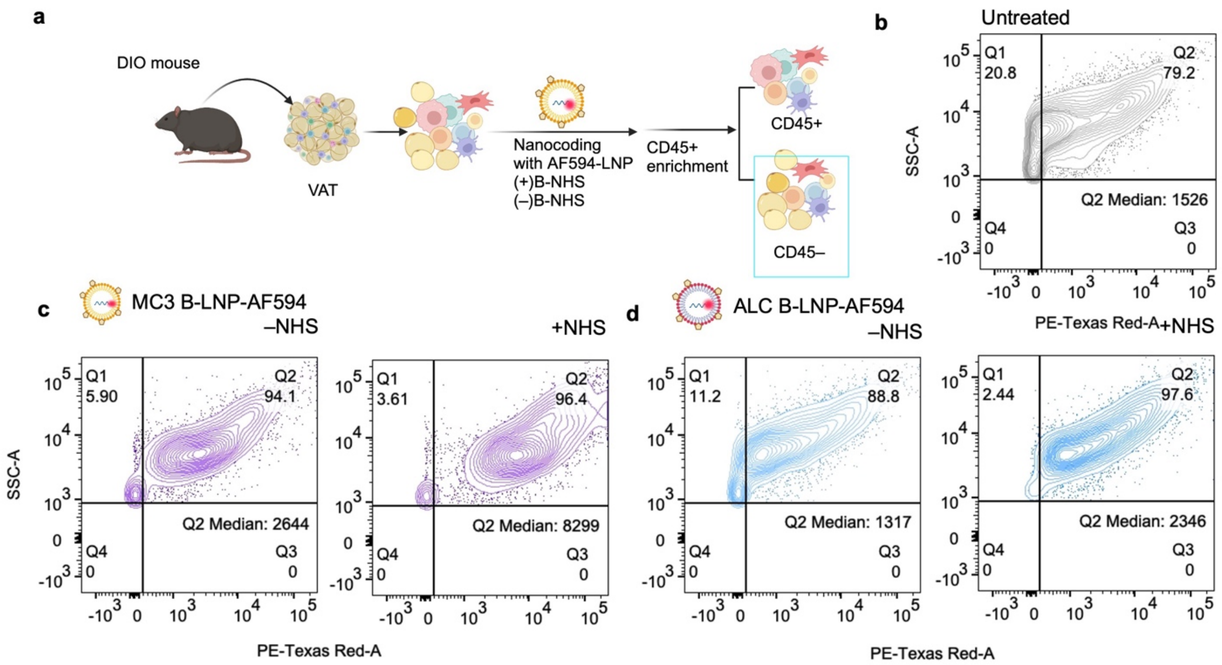

**Figure S16.** B-LNP labeling of CD45⁻-enriched mouse adipose tissue stromal vascular fraction (SVF), comparing LNPs based on MC3 and ALC lipids. **(a)** Schematic depiction of SVF cell collection, enrichment for CD45⁻ cells, and labeling with B-LNP containing AF594-DNA (**Table S1**). Cells were first labeled with B-NHS and SAv (not shown) before the addition of B-LNP Flow cytometry data are shown for CD45⁻ SVF cells, including **(b)** untreated cells to show autofluorescence, **(c)** cells treated with MC3-based B-LNP, and **(d)** cells treated with ALC-based B-LNP. The median AF594 intensity values of cells in Q2 are shown in each figure. Due to the extensive autofluorescence of these cells, labeling is obscured, but cells labeled with ALC-based B-LNPs have median intensity similar to that of the autofluorescence background. The median intensity of cells in Q2 is higher for the MC3-based LNP (8299) *versus* the ALC-based LNP (2346). Each experimental group applied the same gates.

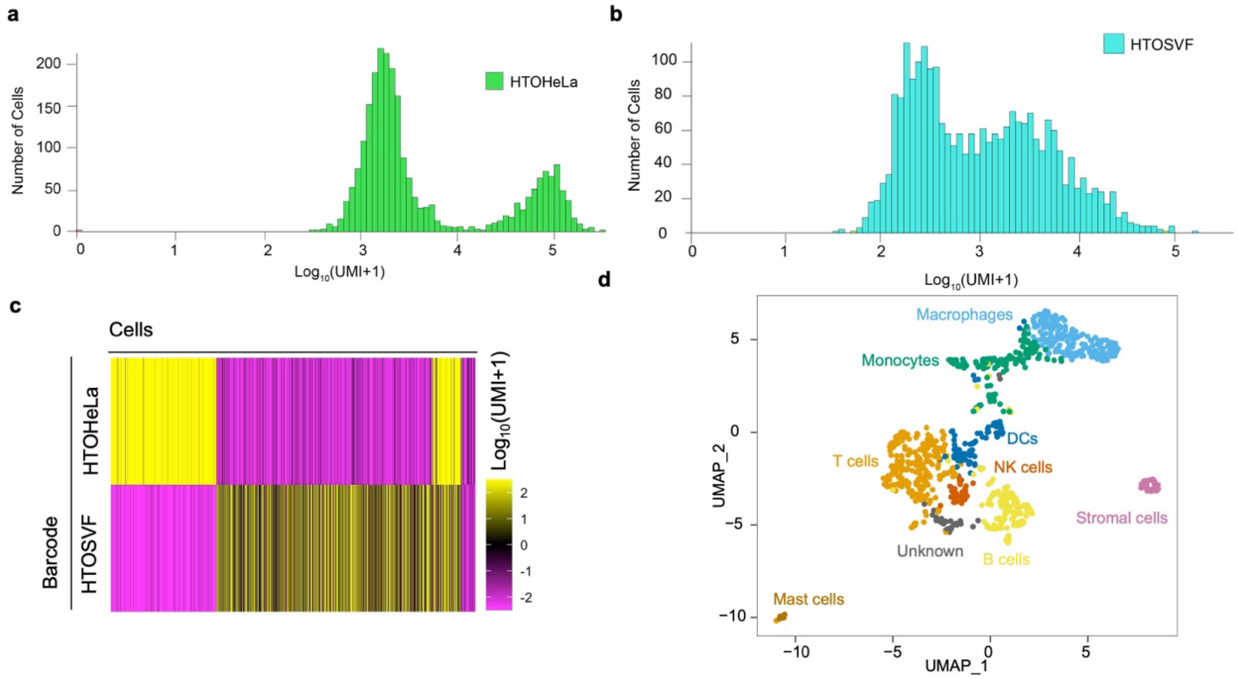

**Figure S17.** Data analysis for the species-mixing study using Nanocoding-based scRNA-seq, using a mixture of mouse SVF cells and human HeLa cells. **(a)** Raw barcode counts for HTO-HeLa. **(b)** Raw barcode counts for HTO-SVF. The histograms were plotted using the CellRanger v7.0 package.[5] **(c)** Heatmap of $\log_{10}$(UMI+1) reads of 2 barcode sequences (2 rows) in cells (thin columns). Cells were sorted by the identified sample origin determined by deMULTIplex package.[2] **(d)** SVF cell types manually annotated in UMAP representation in RNA space.

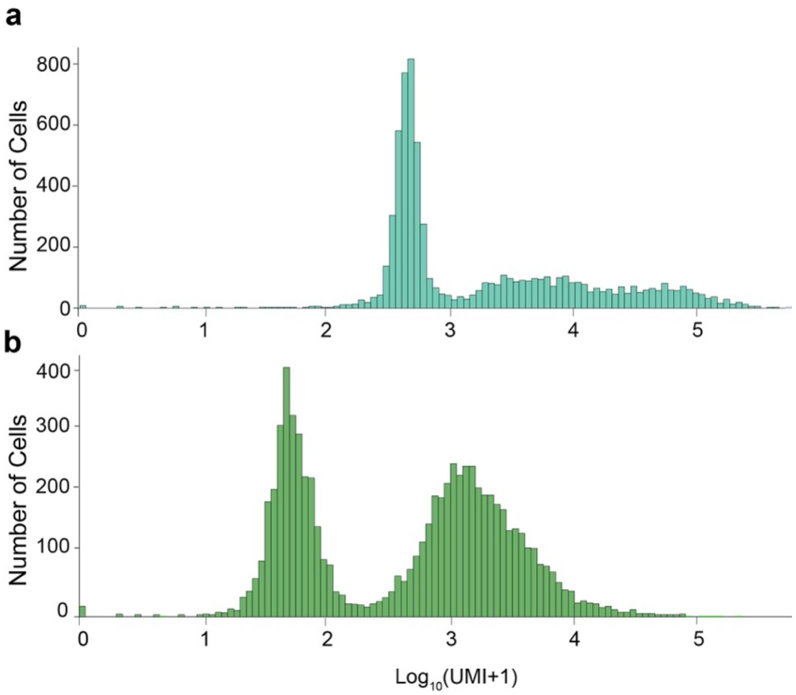

**Figure S18.** Raw barcode count histogram for the young+aged adipose study**. (a)** Raw barcode counts for HTO-Young. **(b)** Raw barcode counts for HTO-Aged. The histograms were plotted using CellRanger v7.0 package.[5]

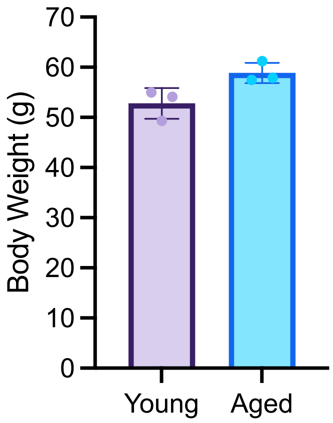

**Figure S19.** Body weight of mice in the young+aged adipose study.

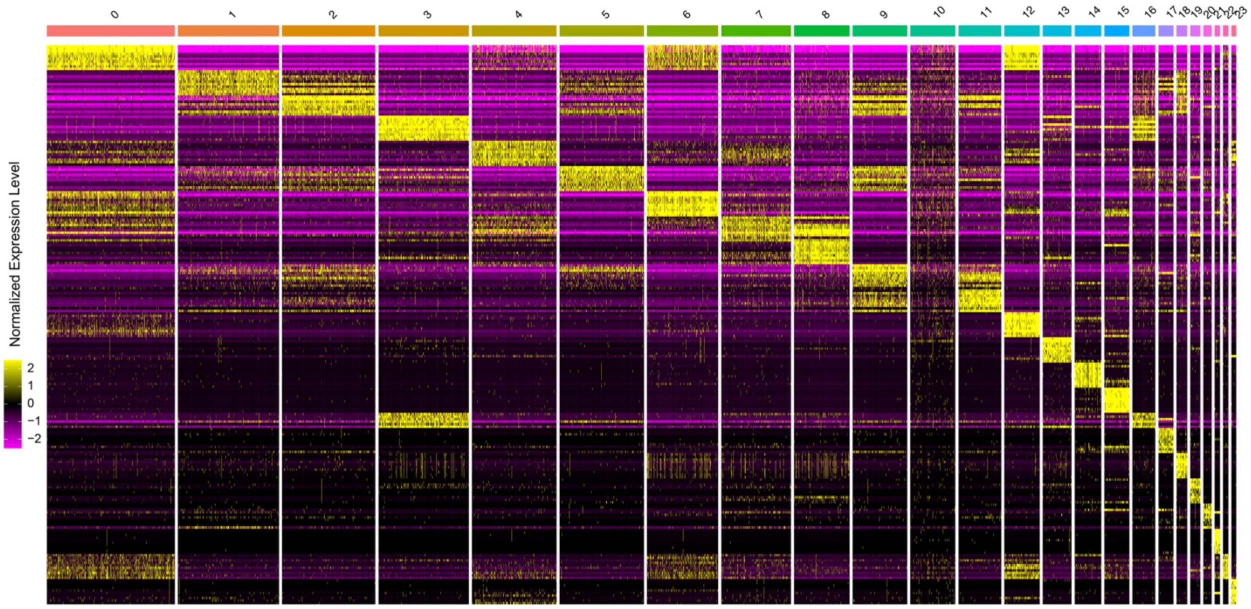

**Figure S20.** Top 10 variated genes for manually annotated cell clusters in the young+aged adipose study. Each row is a unique gene, and each thin column is one cell. Cells are clustered by RNA expression with 24 clusters identified by FindClusters in Seurat (details in **Methods**). The top genes are sorted by fold change for each cluster. The list of top 10 genes in each cluster and corresponding cell type are in **Table S7.** The plot was generated by the Seurat v5 package.

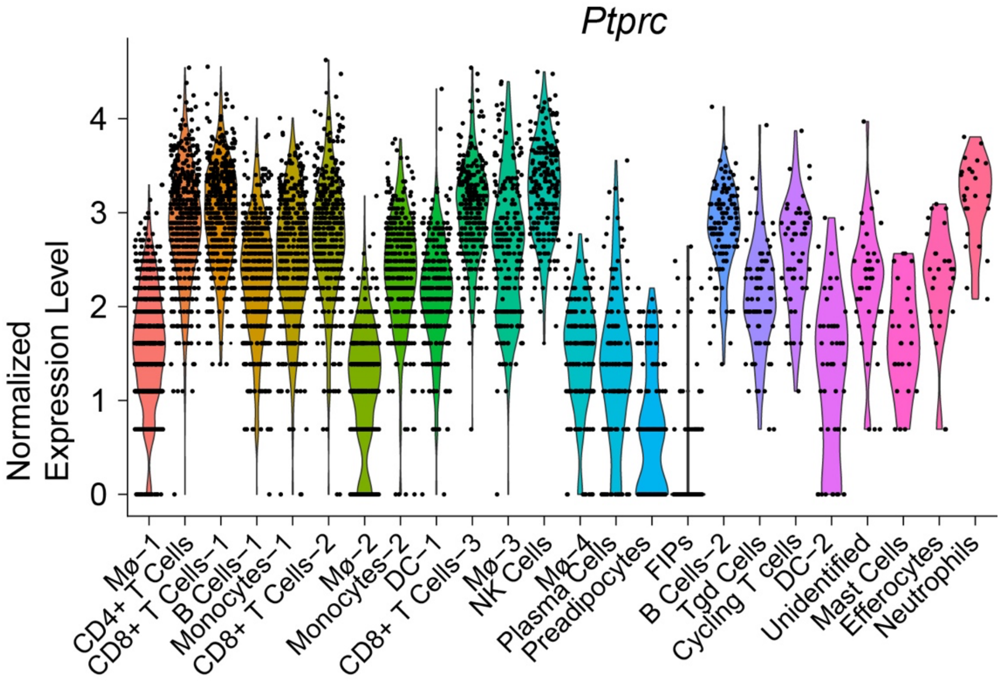

**Figure S21.** Distribution of expression of *Ptprc* (gene encoding CD45 protein) in each cell type identified in the young+aged adipose study.

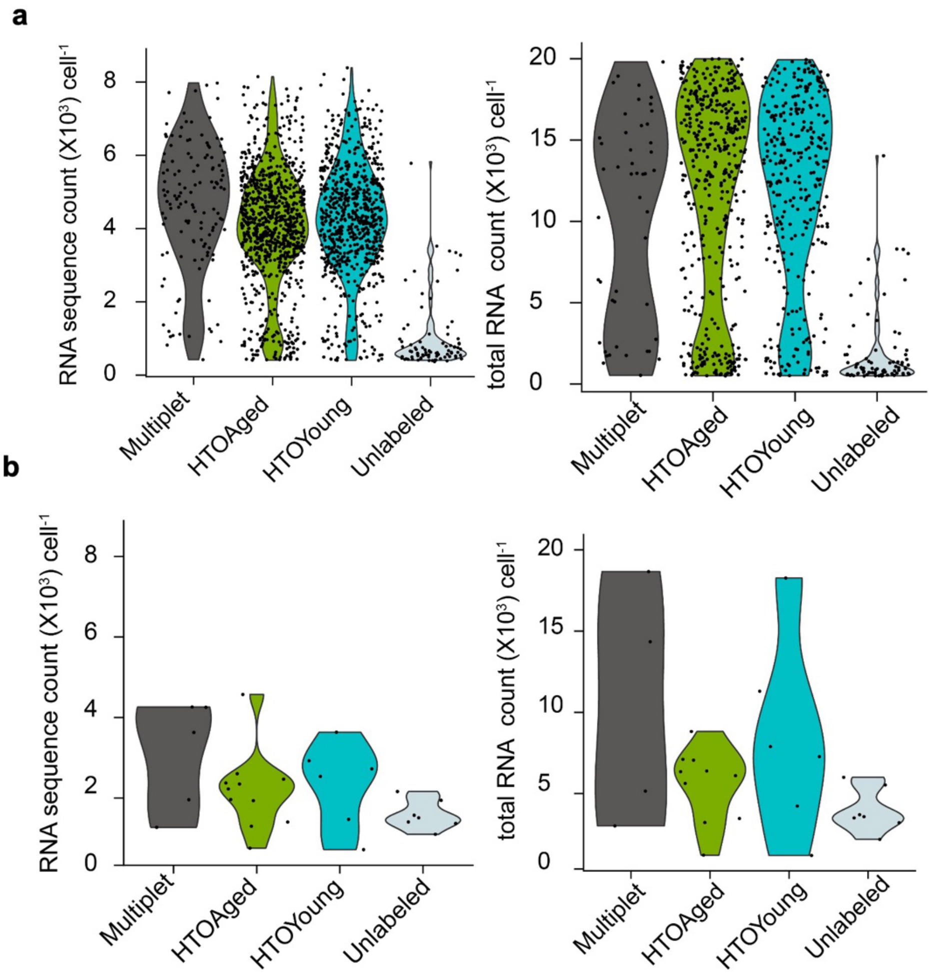

**Figure S22.** RNA quality evaluation of macrophages and mast cells in the young+aged adipose study. Data show **(a)** macrophages and **(b)** mast cells, showing RNA sequence count and total RNA count correspond to nFeature_RNA and nCount_RNA, respectively, from Seurat v5.

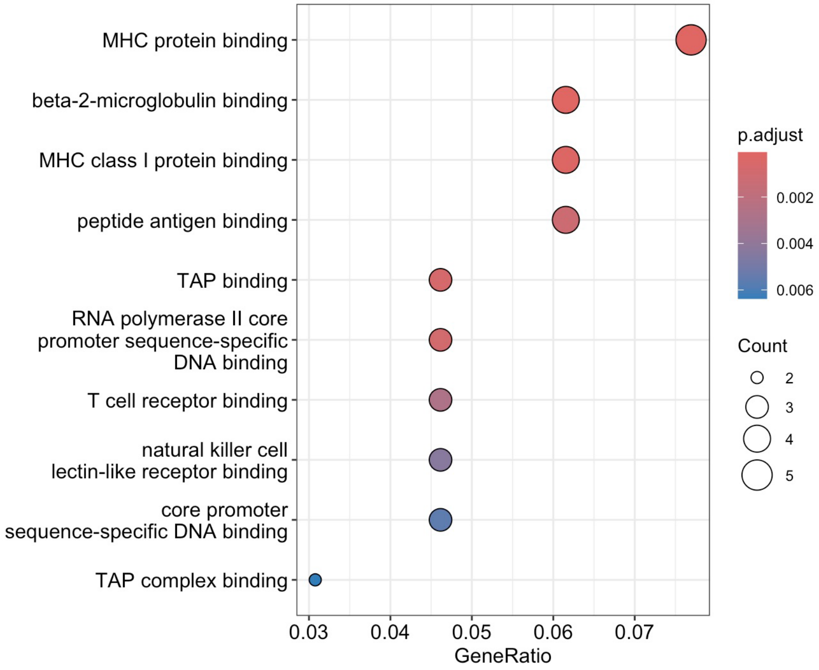

**Figure S23.** Enriched gene ontology (GO) terms for genes upregulated in macrophages in the aged group from the young+aged adipose study. Ten GO molecular functions with the largest gene ratios are plotted. The dot size represents the number of genes in the significant differentially expressed genes (DEG) list associated with the GO term and the color represents the *p*-adjusted value. The GO analysis was conducted using the ClusterProfiler package.[6]

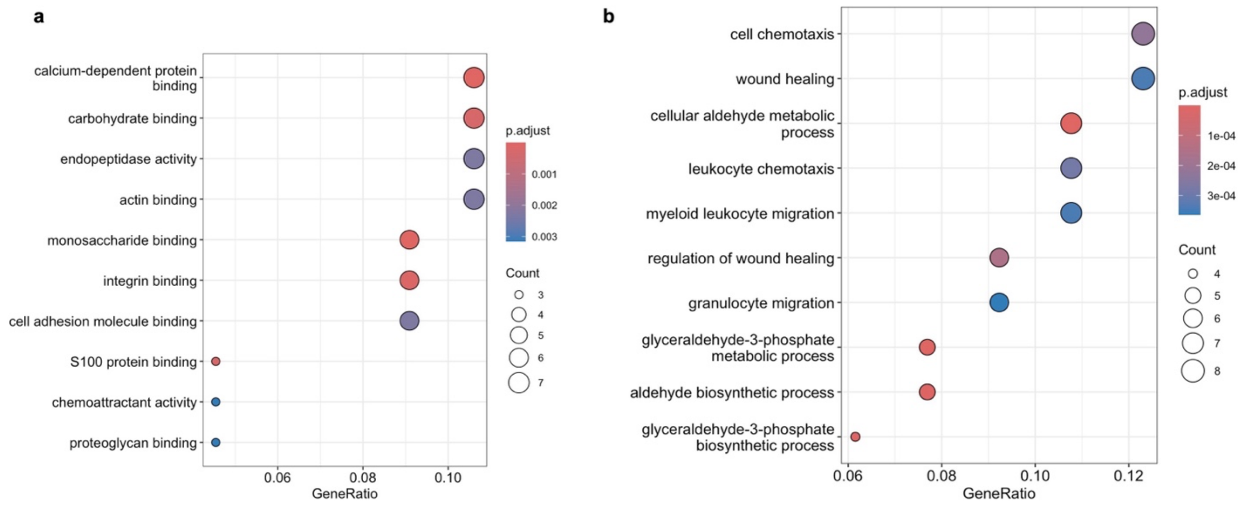

**Figure S24**. Enriched gene ontology (GO) terms for genes upregulated in macrophages in the young group from the young+aged adipose study. Plots show **(a)** ten GO molecular functions with the largest gene ratios and **(b)** ten GO biological processes with the largest gene ratios. The dot size represents the number of genes in the significant DEG list associated with the GO term and the color represent the *p*-adjusted value. The GO analysis was conducted using the ClusterProfiler package.[6]

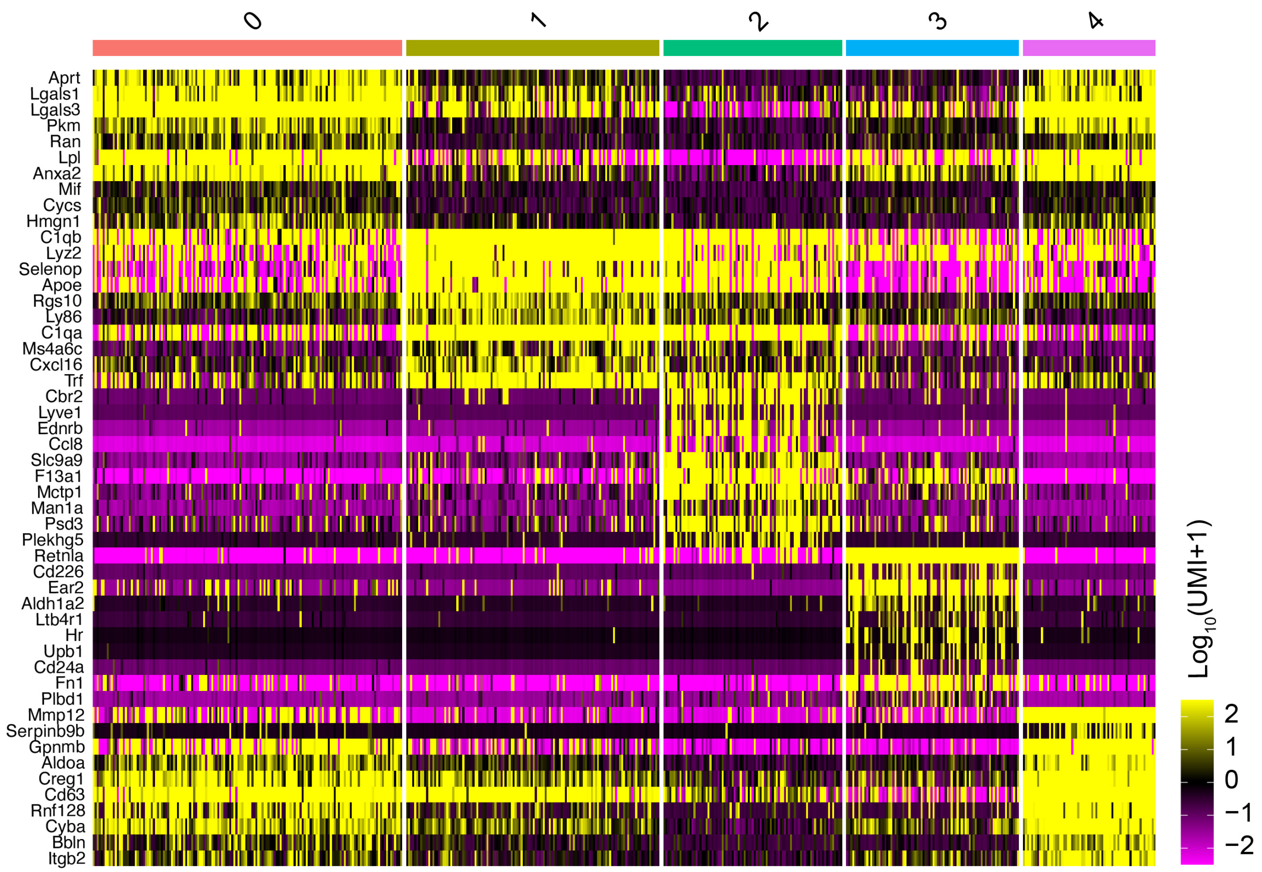

**Figure S25.** Gene expression in macrophage subtypes from the young group in the young+aged adipose study. The top 10 expressed genes are shown in each subtype. The cluster numbers at top refer to the cluster numbers in **Figure 5h**. Each cluster was manually annotated using the marker genes and references in **Table S8**.

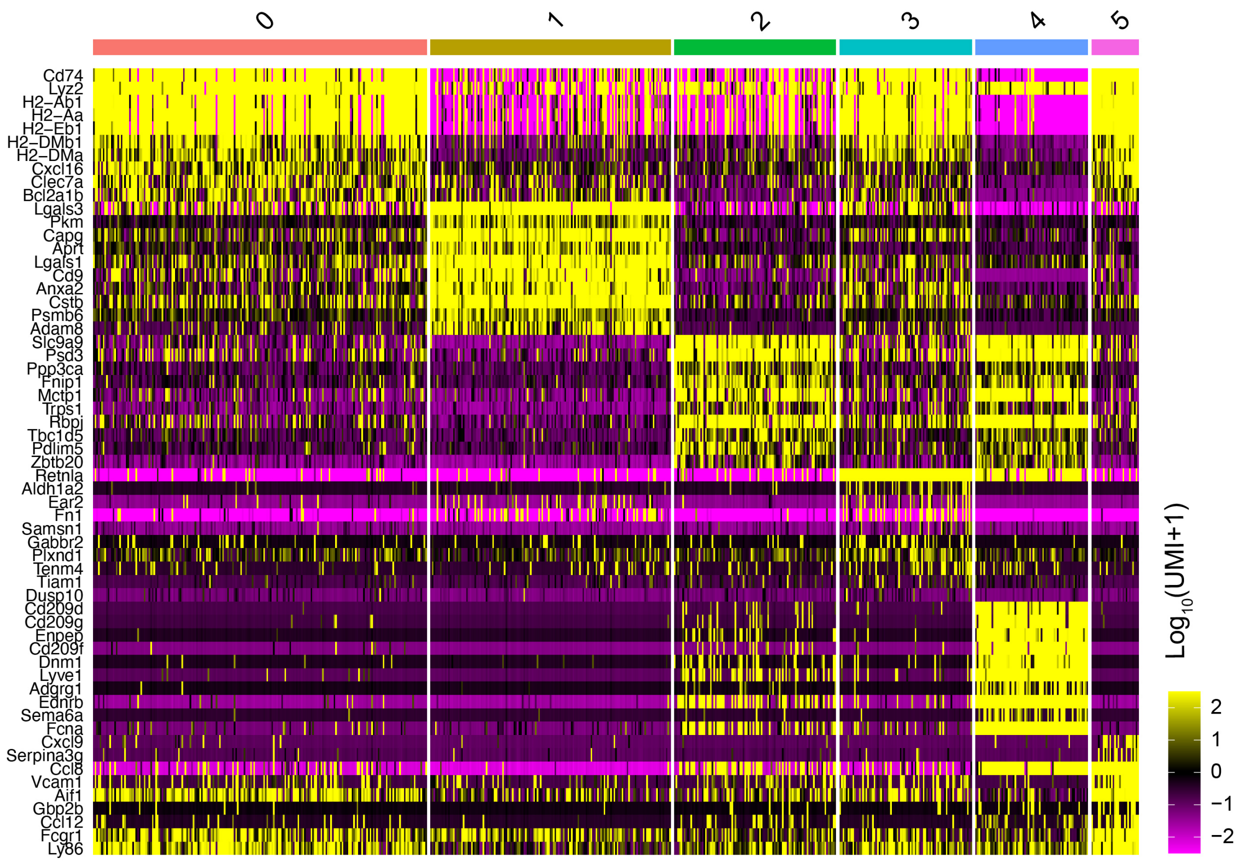

**Figure S26.** Gene expression in macrophage subtypes from the aged group in the young+aged adipose study. The top 10 expressed genes are shown in each subtype. The cluster numbers at top refer to the cluster numbers in **Figure 5i**. Each cluster was manually annotated using the marker genes and references in **Table S8**.

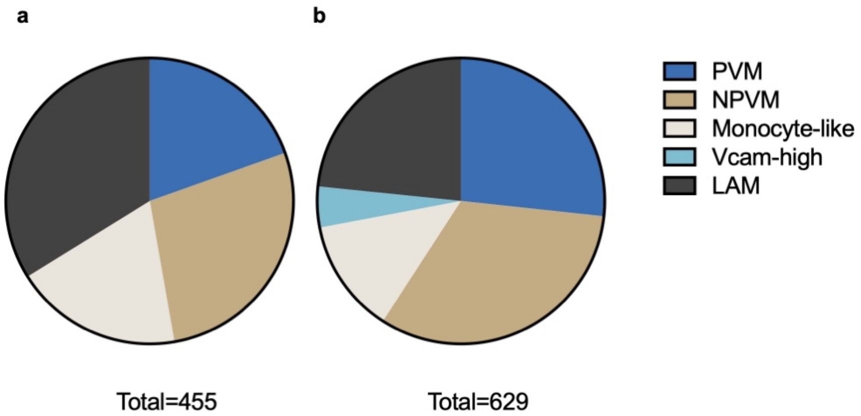

**Figure S27.** Distribution of each macrophage subtype in the young+aged adipose study. Charts show (**a**) young group with clustering as in **Figure S24** and (**b**) aged with clustering as in **Figure S25**.

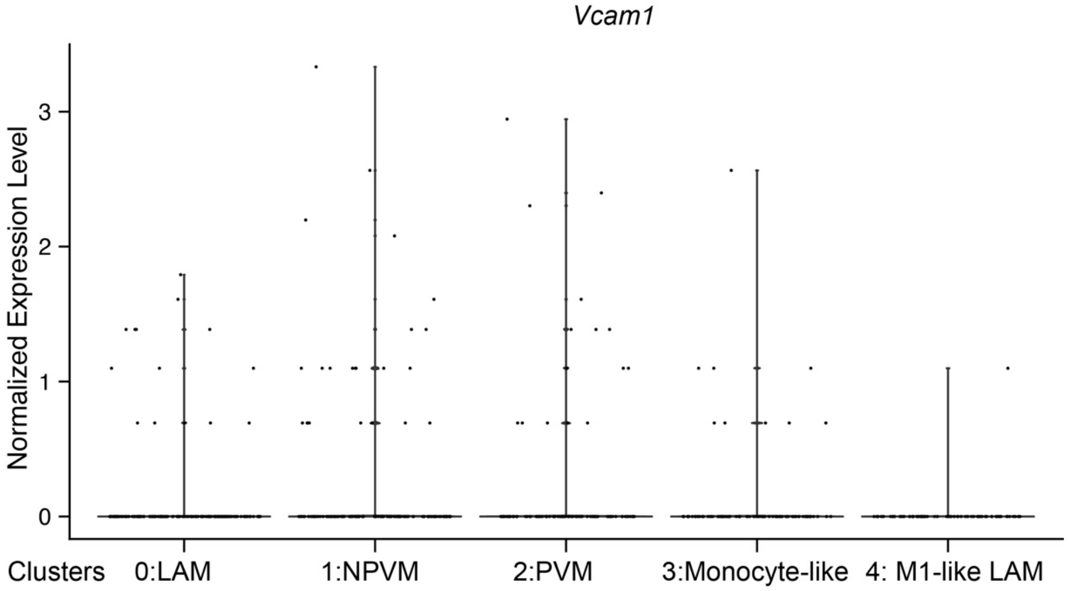

**Figure S28.** *Vcam1* expression in macrophage subtypes in the young group from the young+aged adipose study. The young group macrophages were independently clustered. The cluster numbers are the same as those in **Figure S24** and manually annotated as in **Figure 5h**.

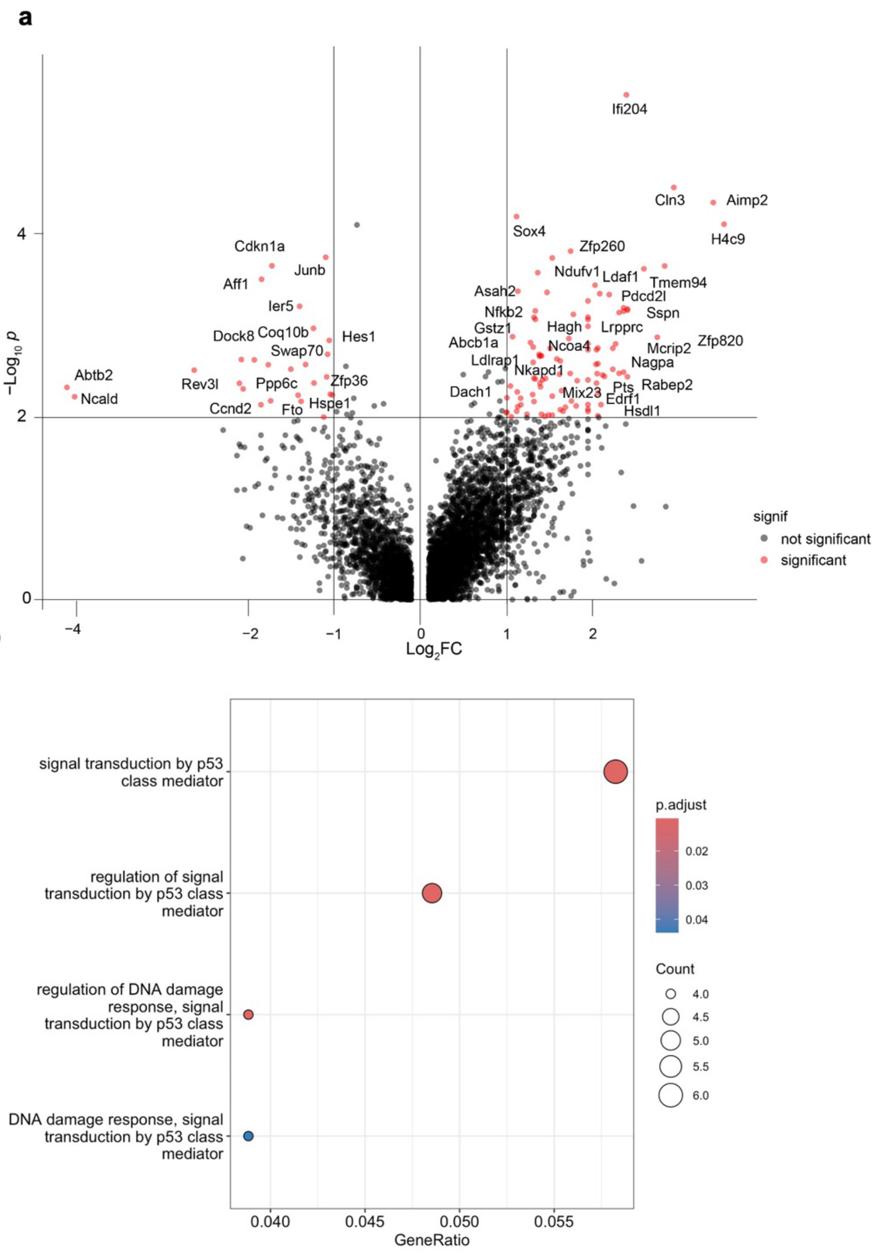

**Figure S29.** Preadipocyte gene expression in the young+aged adipose study. **(a)** Differential gene expression of preadipocytes comparing aged *versus* young. Selected top variated genes are labeled. **(b)** Gene ontology biological process terms of genes upregulated in the aged group. The GO analysis was conducted using the ClusterProfiler package.[6]

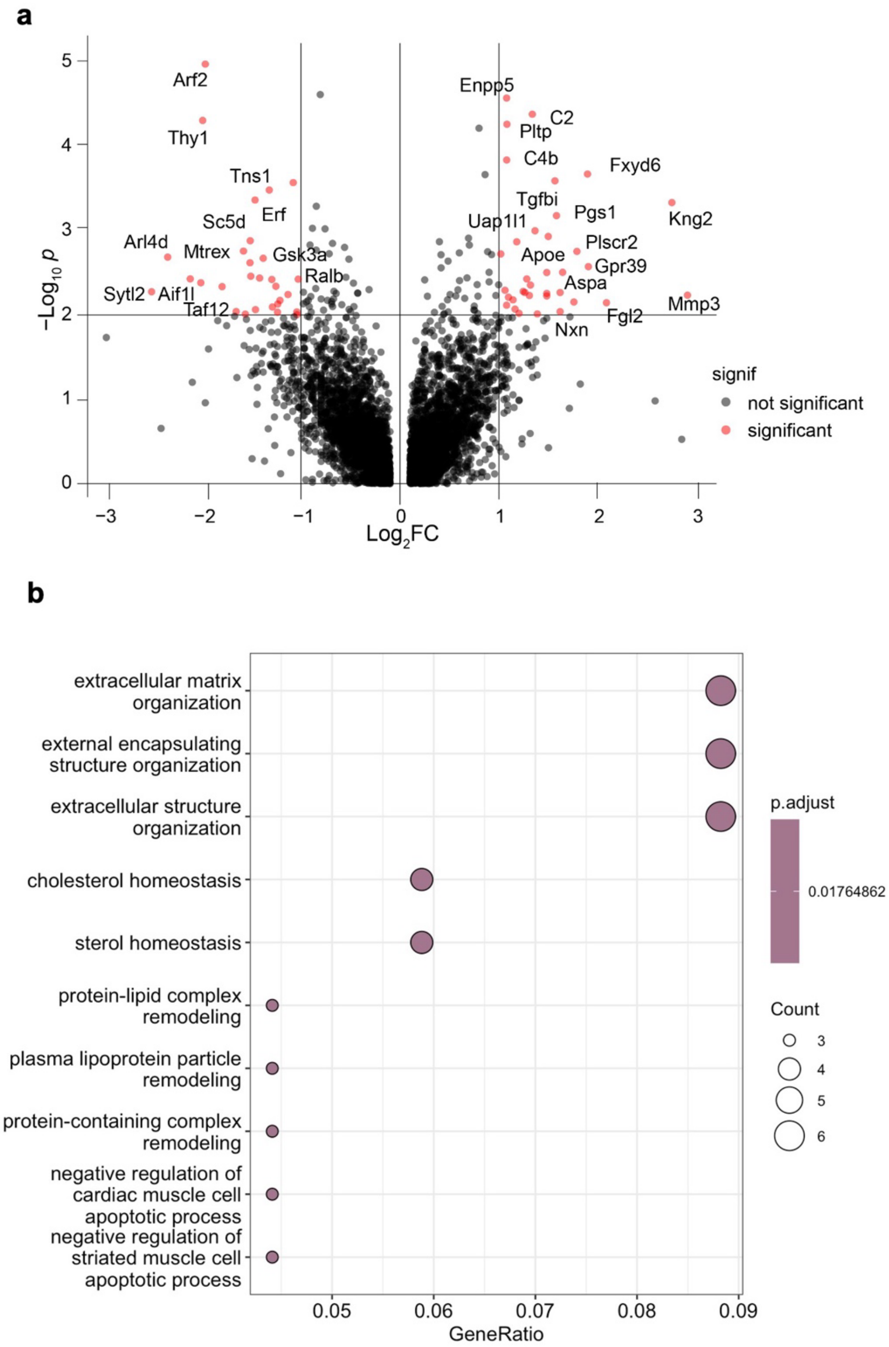

**Figure S30.** Fibro-inflammatory progenitor (FIP) cell gene expression in the young+aged adipose study. **(a)** Differential gene expression of FIP cells comparing aged *versus* young. Selected top variated genes are labeled. **(b)** Gene ontology biological process terms of genes upregulated in the aged group. The GO analysis was conducted using ClusterProfiler package.[6]

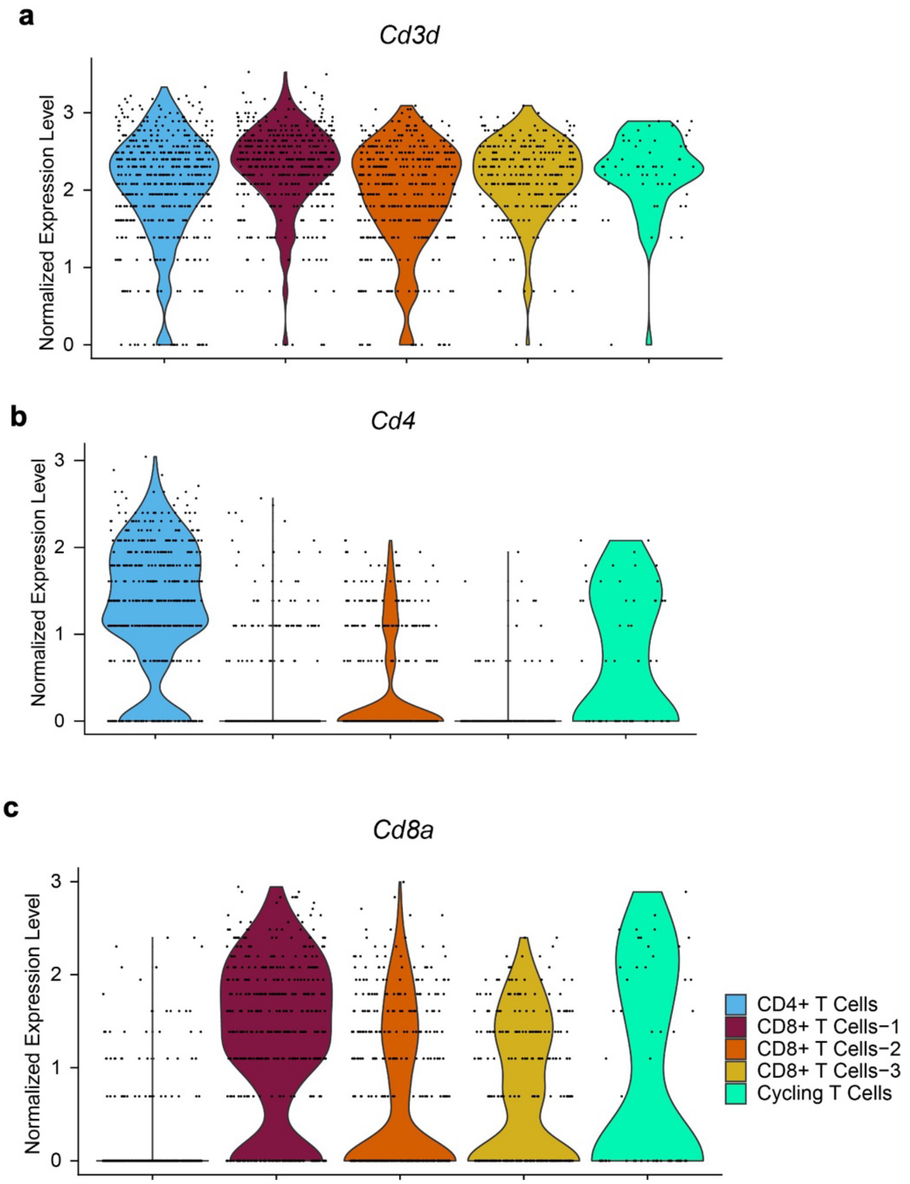

**Figure S31.** Marker genes to determine T cell subtypes in the young+aged adipose study. Expression levels are shown for **(a)** *Cd3d*, **(b)** *Cd4*, and **(c)** *Cd8a* for the young and aged groups together.

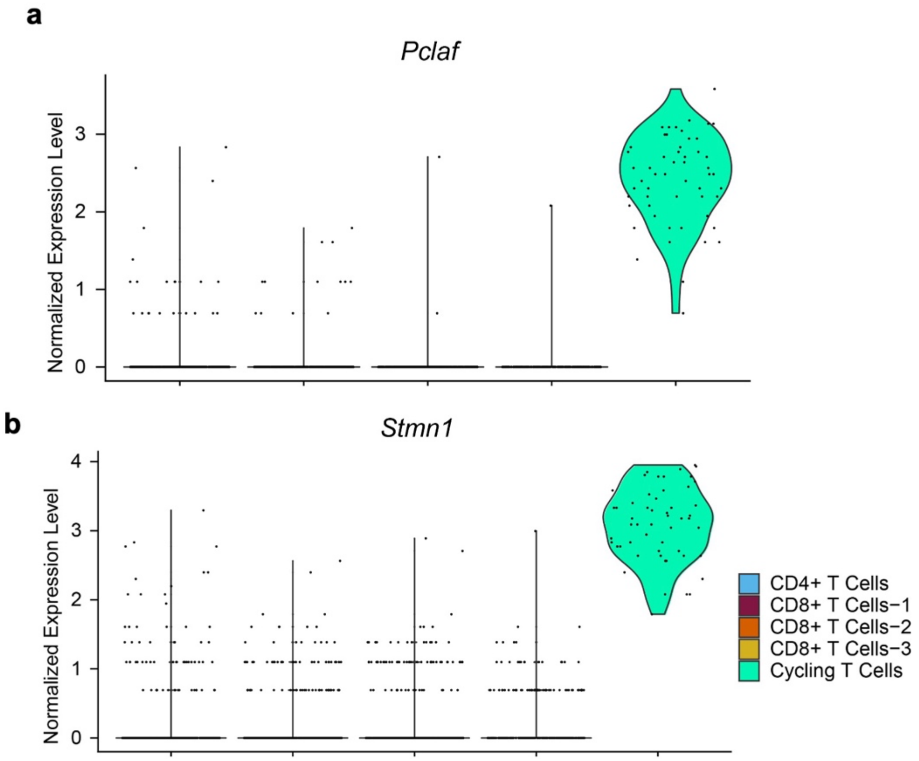

**Figure S32.** Marker genes to determine the ‘cycling T cell’ cluster in the young+aged adipose study. Expression levels of cell cycle marker genes are shown for **(a)** *Stmn1* and **(b)** *Pclaf* for the young and aged groups together. This cluster is defined as cycling T cells based on reported cell annotations[7] and set as the starting point in pseudotime analysis in **Figure 5k**.

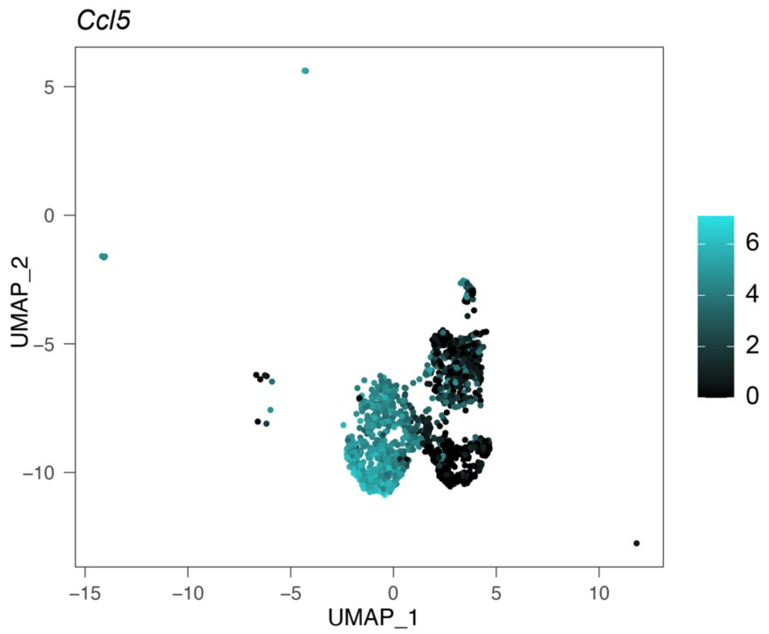

**Figure S33.** *Ccl5* gene expression overlay on the UMAP of T cells in the aged group from the young+aged adipose study. The plot is the uncropped version of **Figure 5m**.

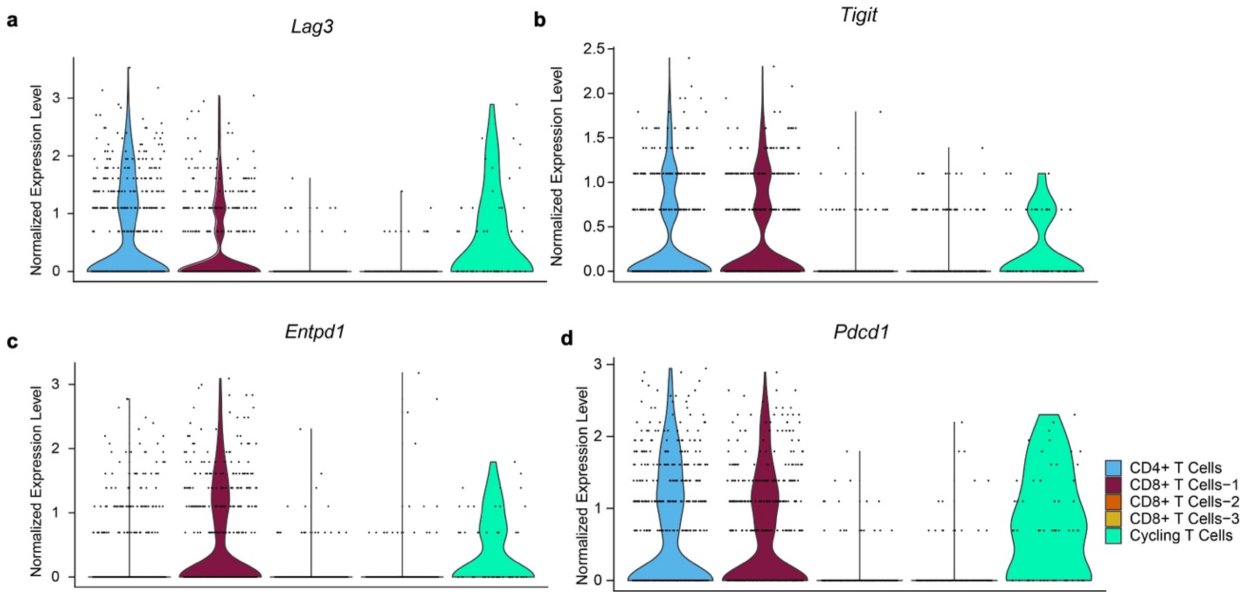

**Figure S34.** Marker genes to determine T cell exhaustion in the aged group in young+aged adipose study. Expression levels are shown for **(a)** *Lag3*, **(b)** *Tigit*, **(c)** *Entpd1*, and **(d)** *Pdcd1* in each T cell cluster.

#### 2 Supporting Tables

##### Table S1. Oligonucleotide sequences

| Application | Oligonucleotide | Sequence |
|---|---|---|
| **Flow cytometry & Fluorescence microscopy** | AlexaFluor488-ssDNA (AF488-DNA) | 5'- /5Alex488N/ CAG ATT AAA CGT GCC ATA CC -3' |
| | AlexaFluor594-ssDNA (AF594-DNA) | 5'- /5Alex594N/ CAG ATT AAA CGT GCC ATA CC -3' |
| | AF488-ssDNA-Biotin (B-DNA-AF488) | 5'- /5Alex488N/ CAG ATT AAA CGT GCC ATA CC /3Bio/ -3' |
| **qPCR** | 55-mer | 5'- /5AmMC6/ GGC AGG AAG ACA AAC ACT GTG CTT GTT ACT GAA CTC TAG TGG TCT GTG GTG CTG T -3' |
| | Biotin-55-mer | 5'- /5Biosg/ GGC AGG AAG ACA AAC ACT GTG CTT GTT ACT GAA CTC TAG TGG TCT GTG GTG CTG T -3' |
| | 55-mer Forward Primer | 5'- GGC AGG AAG ACA AAC A -3' |
| | 55-mer Reverse Primer | 5'- ACA GCA CCA CAG ACC A -3' |
| | 'Spike-in' Forward Strand | 5'- CGC GAG ATA CAC TGC CAG AAA TCC GCG TGA TTA CGA GTC GTG GTA AAT TTA ATC TGG CTG TGG TC -3' |
| | 'Spike-in' Reverse Strand | 5'- GAC CAC AGC CAG ATT AAA TTT ACC ACG ACT CGT AAT CAC GCG GAT TTC TGG CAG TGT ATC TCG CG -3' |
| | 'Spike-in' Forward Primer | 5'- GAC CAC AGC CAG ATT AAA TTT ACC A -3' |
| | 'Spike-in' Reverse Primer | 5'- CGC GAG ATA CAC TGC CAG AA -3' |

**Table S1. (continued)**

| Application | Oligonucleotide+ | Sequence* |
|---|---|---|
| **Sequencing** (90mer ssDNA) | HTO-1 (Obese-1) | 5'- /5AmMC6/ GTG ACT GGA GTT CAG ACG TGT GCT CTT CCG ATC TNN NNN NNN NNA AGG CAG ACG GTG CAN NNN NNN NNG CTT TAA GGC CGG TCC TAG CAA -3' |
| | HTO-2 (Obese-2) (HTO-Young) | 5'- /5AmMC6/ GTG ACT GGA GTT CAG ACG TGT GCT CT CCG A T T N NNN NNN NNG ACG CGC GTT GTC A T NNN NNN NNG CT TAA GGC CGGTCC TAG CAA -3' |
| | HTO-3 (Obese-3) (HTO-HeLa) | 5'- /5AmMC6/ GTG ACT GGA GTT CAG ACG TGT GCT CTT CCG A T T N NNN NNN NNG TGC AAG AGT TGG CGN NNN NNN NNG CTT TAA GGC CGGTCCTAG CAA -3' |
| | HTO-4 (Lean-1) | 5'- /5AmMC6/ GTG ACT GGA GTT CAG ACG TGT GCT CTT CCG A T T N NNN NNN NNG CCA AGA TCA GGT CCNNNN NNN NNG CIT TAA GGC CGGTCCTAG CAA -3' |
| | HTO-5 (Lean-2) | 5'- /5AmMC6/ GTG ACT GGA GTT CAG ACG TGT GCT CT CCG A T T N NNN NNN NNC ACT CCT TGA CAG GIN NNN NNN NNG C T TAA GGC CGGTCC TAG CAA -3' |
| | HTO-6 (Lean-3) (HTO-Aged) | 5'- /5AmMC6/ GTG ACT GGA GTT CAG ACG TGT GCT CTT CCG A T T N NNN NNN NNG CAT TGC GTC AGG C T NNN NNN NNG C T TAA GGC CGG TCC TAG CAA -3' |
| | HTO-SVF | 5'- /5AmMC6/ GTG ACT GGA GTT CAG ACG TGT GCT CTT CCG A T T N NNN NNN NNG GCT GCG CAC CGC CTN NNN NNN NNG CT TAA GGC CGG TCC TAG CAA -3' |

* N denotes a random base.

+ Names in the parentheses are the corresponding barcode names used in figures and the main text.

#### Table S2. Lipid names, structures, and manufacturers

| Lipid name (abbreviation) | Structure | Manufacturer | Concentration* |
|---|---|---|---|
| DLin-MC3-DMA (MC3) | 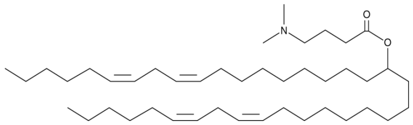 | Cayman Chemical | 10 mg/mL in ethanol |
| ALC-0315 (ALC) | 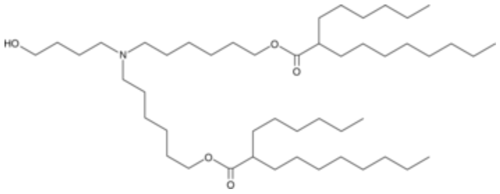 | Cayman Chemical | 10 mg/mL in ethanol |
| SM-102 (SM) | 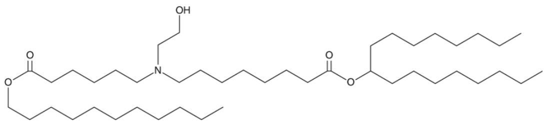 | Cayman Chemical | 10 mg/mL in ethanol |
| DODAB (Non-ionizable lipid, NI) | 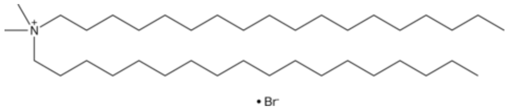 | Cayman Chemical | 10 mg/mL in ethanol |
| Cholesterol | 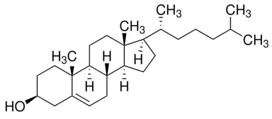 | Sigma Aldrich | 10 mg/mL in ethanol |
| DSPC (18:0/18:0 PC) | 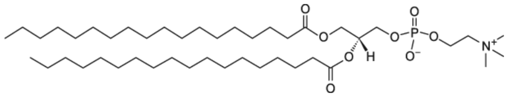 | Avanti Polar Lipids | 10 mg/mL in ethanol |
| DSPE-PEG(2000)-Biotin (B-PEG-DSPE) | 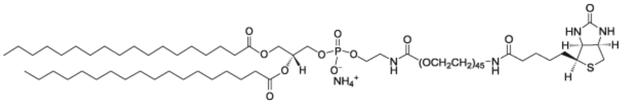 | Avanti Polar Lipids | 10 mg/mL in chloroform |
| DMG-PEG(2000) (PEG-DMG) | 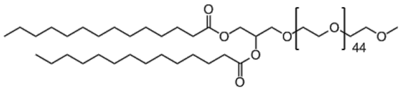 | Avanti Polar Lipids | 10 mg/mL in ethanol |

*This concentration is used for lipid mix without further dilution.

#### Table S3. LNP compositions by mole percentage

| LNP name** | Ionizable or cationic lipid | | Cholesterol | DSPC | PEG-Lipid | | N:P[+] |
|---|---|---|---|---|---|---|---|
| **MC3(B-1.5)-LNP (B-LNP)**[‡] | MC3 | 50.0 % | 38.5 % | 10.0 % | B-PEG-DSPE | 1.5 % | 6:1, 3:1, or 1.5:1 |
| **MC3(B-3)-LNP** | MC3 | 49.3 % | 37.9 % | 9.9 % | B-PEG-DSPE | 2.9 % | 3:1 |
| **MC3(M-1.5)-LNP** | MC3 | 50.0 % | 38.5 % | 10.0 % | M-PEG-DMG | 1.5 % | 3:1 |
| **B-LNP-AF488**[§] | MC3 | 50.0 % | 38.5 % | 10.0 % | B-PEG-DSPE | 1.5 % | 3:1 |
| **B-LNP-AF594**[§] | MC3 | 50.0 % | 38.5 % | 10.0 % | B-PEG-DSPE | 1.5 % | 3:1 |
| **ALC(B-1.5)-LNP** | ALC-0315 | 50.0 % | 38.5 % | 10.0 % | B-PEG-DSPE | 1.5 % | 3:1 |
| **SM(B-1.5)-LNP** | SM-102 | 50.0 % | 38.5 % | 10.0 % | B-PEG-DSPE | 1.5 % | 3:1 |
| **NI(B-1.5)-LNP** | DODAB | 50.0 % | 38.5 % | 10.0 % | B-PEG-DSPE | 1.5 % | 3:1 |

* LNP naming convention corresponds to A(B-#)-LNP where A is the ionizable or cationic lipid type, B is the PEG lipid type, and # is the PEG lipid mole percentage.

[+] Molar ratio of nitrogen of ionizable or cationic lipid to phosphate of ssDNA.

[‡] MC3(B1-.5)-LNP is the formulation used for most cell labeling experiments in this work. B-LNP refers to this formulation if not further specified.

[§] B-LNP recipe encapsulating AF488-DNA or AF594-DNA (**Table S1**) for flow cytometry or microscopy tests.

#### Table S4. Metrics for LNPs encapsulating barcodes (*N*>3, independent samples)

| LNP name | size ± s.d. (nm)* | PDI⁺ | Encapsulation efficiency‡ |
|---|---|---|---|
| **B-LNP** | 100.0 ± 10.8 | 0.11 | 90% |
| **MC3(B-3)-LNP** | 88.3 ± 1.1 | 0.22 | 83% |
| **MC3(M-1.5)-LNP** | 129.6 ± 1.8 | 0.12 | 92% |
| **ALC(B-1.5)-LNP** | 165.2 ± 2.2 | 0.13 | 84% |
| **SM(B-1.5)-LNP** | 148.0 ± 0.8 | 0.12 | 87% |
| **NI(B-1.5)-LNP** | 180.6 ± 0.7 | 0.19 | 85% |

* Hydrodynamic diameter by DLS using Z-average metrics. *N*=6 replicates for each sample.

⁺ Polydispersity index by DLS.

‡ Percentage of encapsulated ssDNA after ultrafiltration to total ssDNA used in each LNP formulation (details in the **Methods**).

#### Table S5. Antibodies and cell labels

| Label | Dilution factor | Manufacturer | Lot # |
|---|---|---|---|
| anti-mouse CD3, PE conjugate | 1:500 | Biolegend | B351058 |
| anti-mouse CD19, FITC conjugate | 1:500 | Biolegend | B369525 |
| anti-mouse CD45, Brilliant Violet 605 conjugate | 1:500 | BD Biosciences | 563053 |
| anti-mouse CD45, AlexaFluor 488 conjugate | 1:500 | Biolegend | B422284 |
| anti-mouse CD45, Brilliant Violet 421 conjugate | 1:500 | Biolegend | 2541104 |
| anti-mouse CD11b, Brilliant Violet 421 conjugate | 1:500 | Invitrogen | B383535 |
| anti-mouse CD11b, AlexaFluor 647 conjugate | 1:500 | BD Biosciences | 1354108 |
| LIVE/DEAD Fixable Near IR (780) Viability Kit for 633 nm excitation | 1:1000 | Invitrogen | 2599157 |
| Vybrant DiO cell-labeling solution | 1:1000 | Invitrogen | 2652728 |
| Cell Mask Orange Plasma membrane stain | 1:1000 | Invitrogen | 2689169 |
| Nuclear Violet LCS1 | 1:1000 | AAT Bioquest | 3240041 |

#### Table S6. scRNA-seq quality reports by CellRanger Count

| Experiment | Suggested kit limit | Input cell number | Cells after Cellranger | Mean reads (cell$^{-1}$) | Median genes (cell$^{-1}$) | Valid UMI (%)[§] |
|---|---|---|---|---|---|---|
| Species mixing study | 1,000* | ~2,000[+]<br>(3:1 SVF:HeLa) | ~2,300[‡] | 522,689 | GRCh:8,078<br>GRCm:3,057 | 95.5% |
| Six-sample spleen study | 10,000 | ~10,000<br>(~1,800 per sample) | 8,199 | 127,968 | 2,597 | 99.8% |
| Young+aged adipose study | 10,000 | ~10,000<br>(1:1 young:aged) | 6,943 | 171,528 | 2,892 | 100.0% |

* Pilot test applied a low-throughput kit.

[+] Extra cells were intentionally loaded to form doublets to evaluate accuracy

[‡] This number is higher than the loaded cell number due to calling of low RNA quality droplets as cells. These were identified and removed in downstream analyses (details in **Code**).

[§] Percentage of UMI (not containing Ns or is homopolymer, etc.) valid for copy number counting.

#### Table S7. Top 10 marker gene list for each cluster*

| Cluster + | Cell Type ‡ | Top 10 Genes § |
|---|---|---|
| 0 | Mø-1 | *C1qb, C1qc, C1qa, Adgre1, C3ar1, Pf4, Fcgr3, Pltp, Ms4a7, Apoe* |
| 1 | CD4+ T Cell | *Cd4, Tnfrsf4, Ctla4, Icos, Izumo1r, Cd3g, Tnfsf8, Cd3e, Tox, Hif1a* |
| 2 | CD8+ T Cell-1 | *Nkg7, Ccl5, Gzmk, Ms4a4b, Ctla2a, Cd8a, Cd3d, Cd3e, Cd3g, Hcst* |
| 3 | B Cell-1 | *Cd79a, Ms4a1, Pax5, Cd79b, Ebf1, Ly6d, Bank1, Fcmr, H2-DMb2, Ralgps2* |
| 4 | Monocytes-1 | *Lst1, Sirpb1c, Cx3cr1, Emilin2, Cybb, Gda, Fgr, Cebpb, Clec4a1, Klra2* |
| 5 | CD8+ T Cells-2 | *Lef1, Satb1, Tcf7, Grap2, Ccr7, Txk, Rpl12, Dapl1, Prkcq, Rps16* |
| 6 | Mø-2 | *Trem2, Lgals3, Atp6v0d2, Rnf128, Gpnmb, Capg, Prdx1, Vat1, Cd63, Anxa1* |
| 7 | Monocytes-2 | *Gm2a, Slamf9, Tnip3, H2-DMb1, Olfm1, Plbd1, H2-Ab1, Lsp1, Cd209a, H2-DMa* |
| 8 | DC-1 | *Clec9a, Ifi205, Septin3, Xcr1, Plbd1, Naga, Flt3, Cd24a, Htr7, Naaa* |
| 9 | CD8+ T Cells-3 | *Ly6c2, Sidt1, Gramd2b, Txk, Il18r1, Klrd1, Ctsw, Samd3, Yes1, Cd7* |
| 10 | Mø-3 | *Aopep, Atxn1, Cdk8, Hip1, Camk4, Pdpk1, Abcb1b, Agfg1, Themis, Ptpn22, Ppp6r3, Vps54* |
| 11 | NK Cells | *Ncr1, Klre1, Klrb1c, Spry2, Xcl1, Klrk1, Il12rb2, Adamts14, Il2rb, Ctsw* |
| 12 | Mø-4 | *Lyve1, Cbr2, Ednrb, Plekhg5, Fcna, Ltc4s, Pdgfc, Gas6, Ccl8, C4b* |
| 13 | Plasma Cells | *Derl3, Jchain, Pls1, Oosp1, Cacna1s, Prg2, Fkbp11, Slpi, Tnfrsf17, Chst1* |
| 14 | Preadipocytes | *Chst1, Cldn5, Cavin2, Egfl7, Ptprb, Gpihbp1, Cdh5, Adgrf5, Tm4sf1, Cav1, Cdh13* |
| 15 | Fibro-inflammatory Progenitors (FIP) | *Tnxb, Entpd2, Bicc1, Lum, Col1a1, Mfap5, Dpep1, Scara5, Plac9, Fbn1* |
| 16 | B Cells-2 | *Pax5, Bank1, Ebf1, Ralgps2, Bcl11a, Scd1, Blk, Aff3, Chst3, Mef2c* |
| 17 | Tgd | *Il23r, Scart1, Actn2, Kcnk1, Ppp1r14c, Ly6g5b, Rbpms2, Il1r1, Zfp462, F2r* |
| 18 | Cycling T cells | *Ccna2, H2ac24, Tpx2, Pclaf, Rrm2, H1f5, Kif11, Shcbp1, Hmmr, Pbk* |
| 19 | DC-2 | *Cacnb3, Nudt17, Mreg, Il4i1, Apol7c, Slco5a1, Tmem150c, Strip2, Ccl22, Adcy6* |
| 20 | Undetermined | *Ccdc184, Dach2, Pard3, Il1rl1, Arg1, Klrg1, Calca, Cntn1, Il5, Areg* |
| 21 | Mast Cells | *Mcpt4, Cpa3, Mrgprb1, Fcer1a, Tpsb2, Ms4a2, Mrgprb2, Tph1, Mrgprx2, Hs6st2* |
| 22 | Efferocytes | *Plxna4, Fat3, Slc41a2, St18, Adcy9, Tanc2, Arhgap22, Snx24, Fmn1, Mitf* |
| 23 | Neutrophils | *S100a9, Retnlg, Cxcr2, Il36g, S100a8, Hdc, Trem1, Mmp9, Acod1 Trem3* |

+ The cluster number corresponds to labels in **Figure S20**.

‡ The cell type in most clusters is determined using the Panglao database, which contains gene-cell type associations for scRNA-seq data.[8] FIP, Tgd, cycling T cells, efferocytes, and preadipocytes were determined based on marker genes reported in recent publications listed in **Table S8**.

§ Top variated genes in each cluster are sorted by fold change values, filtered for $p < 0.01$.

**Table S8. Subtype annotation markers for stromal and lymphoid cells in mouse SVF cells in Figure 5c and S20**

| Cluster | Cell subtype | Genes for subtype determination | References |
|---|---|---|---|
| Stromal cells | | | |
| 14 | Preadipocytes | *Car3, Cd36, Fabp4* | Hepler *et al.*[9] |
| 15 | Fibrosis inflammatory progenitor (FIP) | *Ly6c1, Efhd1, Dact2* | Hepler *et al.*[9] |
| Not observed | Mesothelial-like cells | *Upk3b, Pkhd11b, Krt18* | Hepler *et al.*[9] |
| T cells | | | |
| 17 | Tgd cells | *Cd3, Rorc, Il17a*<br>*Cd4* negative, *Cd8* negative | Jensen *et al.*[10] |
| Macrophages | | | |
| 22 | Efferocyte (or metabolically activated macrophage, MME) | *Mertk, Cd36, Abca1* | De Couto *et al*. [11]<br>Yvan-Charvet *et al*. [12] |

#### Table S9. Subtype annotation markers for high-resolution-clustered macrophages in Figure 5h, 5i, and S24-S28

| Cluster | Macrophage subtype | Genes for subtype determination | References |
|---|---|---|---|
| Subsets from young group (**Figure 5h**) | | | |
| 0 | LAM: lipid associated macrophage | *Trem2, Cd9, Lpl, Lipa, Spp1* | Cottam *et al.*[7] Maniyadath *et al.*[13] Jaitin *et al.*[14] Sárvári *et al.*[15] |
| 1 | Non-perivascular macrophage (NPVM)* | *Cd74, Cd14, Clec7a, Il1b, Ccl3* | |
| 2 | Perivascular macrophage (PVM) | *Klf4, Cbr2, Stab1, Mrc1, Cd163, Lyve1* | |
| 3 | Monocyte-like | *Cd74, Rentla, Fn1, Ear2* | |
| 4 | M1-like LAM | *Itgax* (Cd11c encoding gene) and other LAM genes | |
| Subsets from aged group (**Figure 5i**) | | | |
| 0 | NPVM | *Cd74, Cd14, Clec7a ,Il1b, Ccl3* | Cottam *et al.*[7] Maniyadath *et al.*[13] Jaitin *et al.*[14] Sárvári *et al.*[15] |
| 1 | LAM | *Trem2, Cd9, Lpl ,Lipa, Spp1* | |
| 2 | PVM | *Klf4, Cbr2, Stab1, Mrc1, Cd163, Lyve1* | |
| 3 | Monocyte-like | *Cd74*, *Rentla, Fn1, Ear2* | |
| 4 | M2-like PVM | *Clec10a, Mrc1* and other PVM genes | |
| 5 | Not reported | *Vcam1* | |

*The non-perivascular macrophage type showed strong pro-inflammatory features with high expression of cytokines or chemokine signaling and phagocytosis, which was also reported by Sárvári *et al.*[15]
